# A histone hitchhiking mechanism for chromatin maintenance during mammalian transcription elongation

**DOI:** 10.64898/2026.11.16.751813

**Authors:** Yukun Wang, Xizi Chen, Yiqing Luo, John W. Watters, Aixia Song, Fei Xavier Chen, Yanhui Xu, Shixin Liu

## Abstract

Eukaryotic transcription machinery faces the dual challenge of having to dismantle the nucleosome barrier for efficient elongation and faithfully restore the chromatin landscape in its wake to prevent aberrant transcription. The sequence of molecular events from nucleosome unwrapping to reassembly has yet to be delineated. Here, we use a single-molecule platform to visualize the dynamics of chromatin transcription by mammalian elongation complex (EC) in real time. When supplemented with a set of elongation factors and histone chaperones, the EC can efficiently traverse the nucleosome it encounters and, surprisingly, carry the dislodged histones while continuing translocation on DNA, a phenomenon we termed histone “hitchhiking”. The formation of this hitchhiking complex is facilitated by the elongation factor SPT6 and IWS1 through distinct mechanisms. On the other hand, the histone chaperones FACT and NAP1L1 promote the transition from histone hitchhiking to nucleosome reassembly. We further showed that the next nucleosome presents a strong barrier against the hitchhiking complex, thus limiting long-range chromatin disruption. Overall, our results revealed that histone hitchhiking by EC is a prevalent intermediate during chromatin transcription, and the timing and location of its resolution are tuned by multiple factors. This mechanism may provide the robustness and flexibility needed to maintain the local organization of transcriptionally active chromatin.

## Introduction

The eukaryotic chromatin is predominantly organized into nucleosomes, each containing ∼147 base pairs (bp) of DNA wrapped around a histone octamer consisting of one (H3-H4)_2 t_etramer and two H2A-H2B dimers^1^. To transcribe the genomic DNA into messenger RNAs, the transcription machinery—assembled around the RNA polymerase II (Pol II)—is required to efficiently unravel the nucleosomes and displace the histones from the DNA^2,3^. Concurrent with crossing the nucleosome barrier, Pol II and its cofactors must retain the displaced histones and reassemble them into nucleosomes, which are critical for preserving the local chromatin landscape and limiting cryptic transcription^4,5^.

A multitude of elongation factors and histone chaperones have been identified to assist in Pol II transcription through nucleosomes^6–8^. Through elegant biochemical reconstitution, previous studies have made seminal contributions to our understanding of the molecular mechanism of chromatin transcription^9–11^. Recent cryo-electron microscopy studies have provided high-resolution structural snapshots of the EC-nucleosome complex at distinct stages of nucleosome progression^12–17^. Nonetheless, a dynamic view of the chromatin transcription process in an in vitro reconstituted setting has been lacking, particularly for higher eukaryotes. As a result, the order of events during the transition from EC-nucleosome encounter to nucleosome reassembly remains to be determined.

By recording the real-time trajectories of individual molecules, single-molecule techniques have proven to be powerful tools for studying dynamic processes such as transcription and have been employed to characterize Pol II behavior on chromatin templates, mostly in the yeast system^18–20^. Recently, we reported an experimental platform combining optical tweezers and single-molecule fluorescence microscopy to study the kinetic control of mammalian transcription elongation complex (EC) on bare DNA^21^. In the current work, we further developed this assay to directly visualize mammalian transcription elongation on chromatinized DNA. By simultaneously following the fate of interacting EC and nucleosomes over long distances, we identified a prevalent state during chromatin transcription in which the displaced histones remain physically associated with the EC while continuing translocation. We further defined the factors and conditions that facilitate the formation and resolution of this state. These real-time observations provided new insights into how the nucleosome landscape within gene bodies is maintained upon transcription.

### Single-molecule visualization of EC-nucleosome encounters

We reconstituted EC using purified mammalian Pol II and a template/non-template DNA scaffold with a Cy3-labeled RNA primer. The assembly was positioned at the center of a biotinylated DNA template tethered between a pair of optically trapped beads (**Fig. 1a**). The bead-EC assembly was moved to a new channel containing LD655-labeled histone octamers (labeled at histone H4 unless noted otherwise) and the yeast histone chaperone Nap1 (yNap1), which typically yielded 3–7 sparsely loaded nucleosomes on the DNA tether^22^. The chromatinized tether was then transferred to another channel containing ribonucleotides (rNTPs) and purified human elongation factors and histone chaperones (namely, DSIF, TFIIS, SPT6, IWS1, PAF1C, RTF1, ELOF1, P-TEFb, FACT and NAP1L1) to restart transcription (**Fig. S1**). Active complexes were indicated by unidirectional movement of the Cy3 fluorescence signal that marks the EC position. To boost fluorescence signals for the EC, the DNA template immediately downstream of the starting EC position contains seven repeat sequences that, when transcribed into nascent RNA, could each be hybridized to an AlexaFluor488 (AF488)-labeled DNA probe. We used three-color imaging to simultaneously track the behavior of EC and the nucleosomes it encounters during transcription elongation (**Fig. 1b-f**; blue/green for the EC and magenta for the nucleosome).

**Figure 1.**
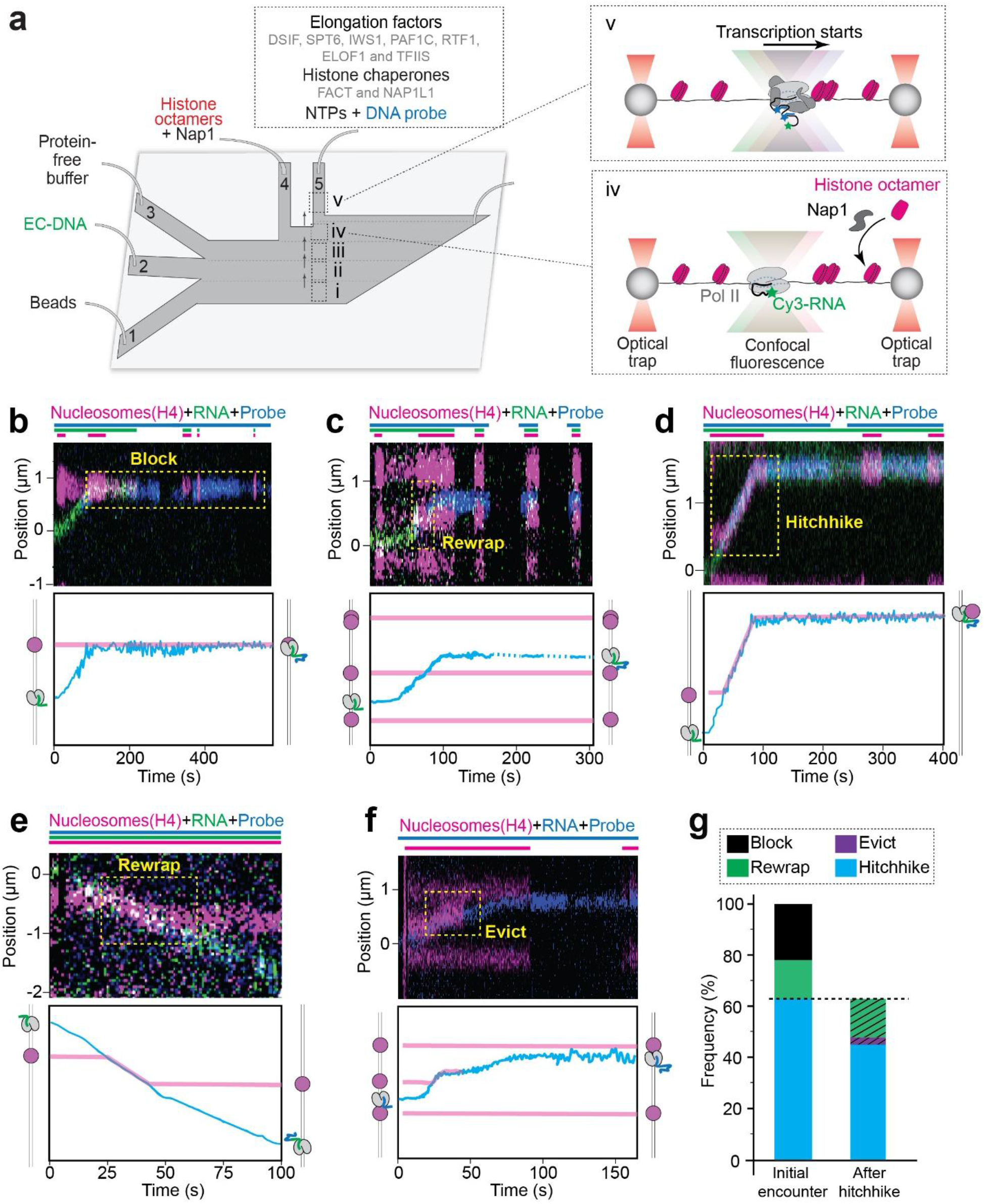
Single-molecule observation of Pol II-nucleosome encounters in a mammalian system. **a**, Schematic of the experimental platform for monitoring mammalian Pol II elongation through nucleosome-loaded DNA. LD655-labeled histone octamers were assembled on tethered DNA with Nap1, followed by transfer to a transcription channel containing NTPs, elongation factors, and histone chaperones. Elongation complexes (ECs) were tracked by labeled RNA and/or DNA probes that hybridize to the nascent RNA. **b-f**, Representative kymographs showing the different outcomes of an EC-nucleosome encounter: long-lived blocking of EC by the nucleosome (**b**), nucleosome rewrapping at the original site after EC passage (**c**), histone hitchhiking with the moving EC (**d**), nucleosome rewrapping at a downstream site (**e**), and histone eviction during hitchhiking (**f**). Bars above the kymographs indicate the time periods when specific lasers were switched on or off, which was intended to distinguish each fluorescence signal and reduce photobleaching. Plots below the kymographs show the extracted EC trajectory (cyan) and histone positions (magenta). **g**, Frequency of each outcome upon initial EC-nucleosome encounter (left). The hitchhiking group is further categorized based on the histone fate at the end of imaging (right). All results in this figure were obtained under the All-Factor condition.

When all the factors were present (hereafter referred to as “All-Factor” condition), we found that the vast majority (∼80%) of ECs traversed the first nucleosome either immediately or after a brief pause (average pause time = 14 ± 3 s, mean ± s.e.m.) (**Fig. S2a**), while only ∼20% of the ECs were blocked by the nucleosome (defined as those that stalled for at least 300 s without progression) (**Fig. 1b**). The distribution between blocked and traversing ECs did not change significantly when the nucleosomes were loaded by the human ortholog of yNap1 (NAP1L1) or via salt dialysis (**Fig. S2b**), nor did it depend on which histone protein was labeled, as a similar traversal ratio was obtained using fluorescently labeled histone H2B instead of H4 (**Fig. S2c**). These results indicate that, under our reconstituted conditions, a nucleosome does not present a strong barrier to the mammalian EC when the tested regulatory factors are present.

### Nucleosomes display multiple fates upon EC passage

Among the events where the EC successfully passed the nucleosome, we observed two distinct scenarios for the histone fate upon initial EC-nucleosome encounter. In ∼20% of cases (16/83 events), the H4 signal was retained at the same location on DNA after the EC passed the nucleosome and continued moving downstream (**Fig. 1c**), indicating histone redeposition onto the DNA. To evaluate whether these events yielded wrapped histones, we pulled on the same chromatin tether at the end of imaging and found the force-extension curve to exhibit transitions characteristic of a nucleosome or a hexasome (**Fig. S3**)^23^. To our surprise, for the majority of events (∼80%) where EC traversed a nucleosome (67/83), the histone signal did not reappear at the same DNA site or disappear, instead co-traveled with the EC along the DNA (**Fig. 1d**). The occurrence of this phenomenon, which we termed as histone “hitchhiking,” does not depend on the nucleosome loading method (**Fig. S2b**). If unimpeded by obstacles such as an additional nucleosome, the hitchhiking complex (EC + histones) can translocate on DNA for up to several kilobases. The hitchhiking histones can sometimes be deposited onto a downstream DNA site (16/67 events; **Fig. 1e**) or, at a much lower probability, be evicted from the DNA into solution as indicated by disappearance of the histone fluorescence signal (3/67 events; **Fig. 1f**). These observations thus identify the hitchhiking complex as a prevalent species in our reconstituted chromatin transcription system (**Fig. 1g**), in which the EC continues to translocate on DNA while carrying the displaced histones.

### SPT6 and IWS1 play opposing roles in histone retention on EC

Next, we asked which component(s) in the EC are required for the newly identified hitchhiking activity. To this end, we conducted single-molecule experiments in which individual elongation factors were omitted from the reaction mix. We found that the omission of SPT6 (ΔSPT6) led to less efficient EC traversal across the nucleosome (**Fig. S2a and S2c**). Moreover, ΔSPT6 resulted in a drastically increased probability of histone eviction at the initial encounter compared to the All-Factor condition (**Fig. 2a, 2b, S4a, and S4b**). ΔSPT6 also significantly increased the probability of histone loss for the hitchhiking complex and reduced the histone residence time on EC (**Fig. 2a and 2c**). These results suggest SPT6 plays a major role in retaining histones on EC, consistent with its well-established histone chaperone role^24^.

**Figure 2.**
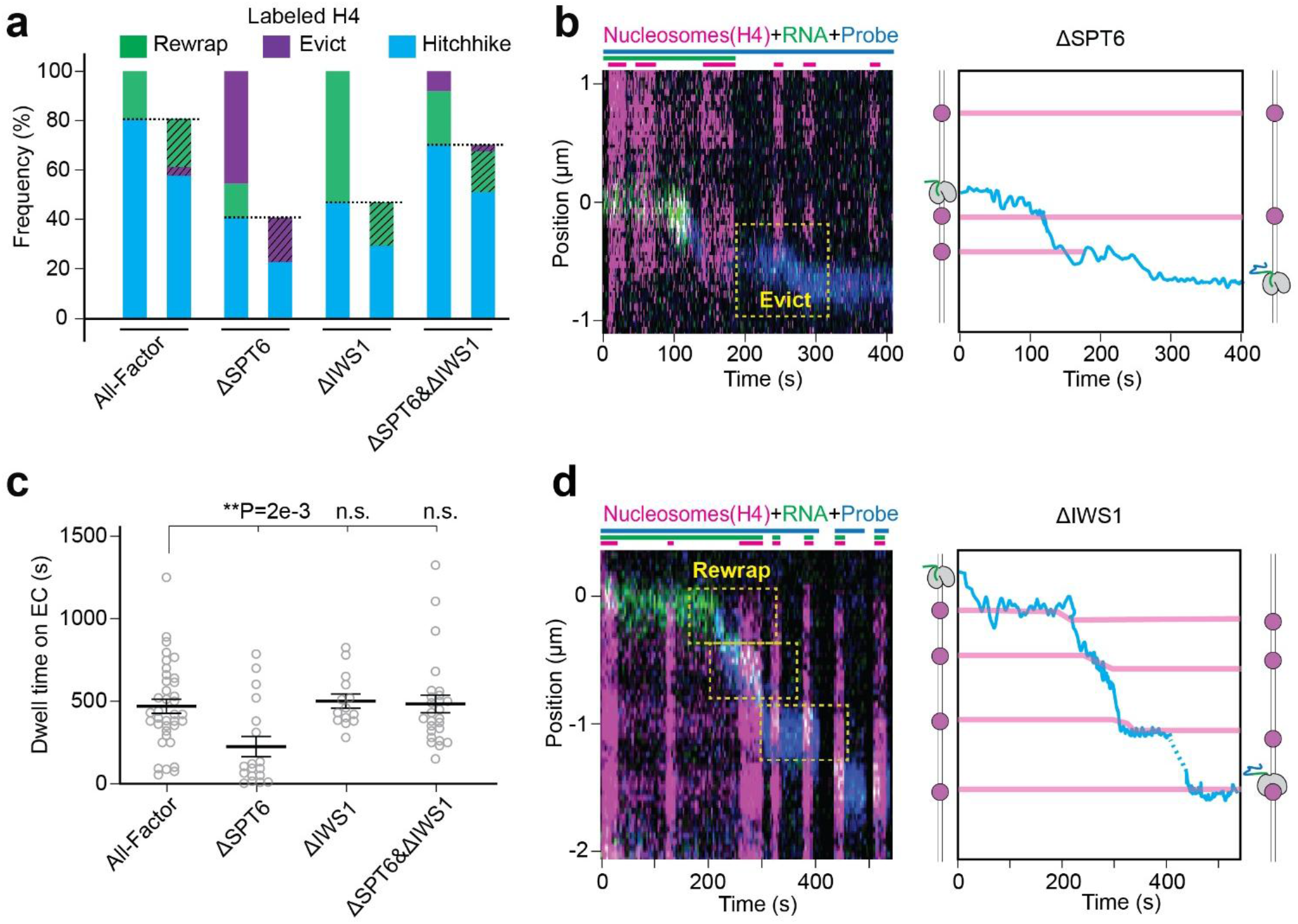
SPT6 and IWS1 play major roles in histone fate determination. **a**, Frequency of different histone H4 fates after an EC passed a nucleosome under the indicated factor conditions. For each condition, both the initial histone fate upon EC-nucleosome encounter (left) and the final histone fate for a hitchhiking complex at the end of imaging (right) are shown. **b**, (Left) Representative kymograph for the ΔSPT6 condition showing the eviction of H4 (loss of its fluorescence signal) after EC passage (left). Bars above the kymograph indicate the time periods when specific lasers were switched on or off. (Right) EC trajectory (cyan) and histone positions (magenta) are extracted from the kymograph to assist with visualization. **c**, Residence time of H4 on EC during hitchhiking under the indicated factor conditions. Each point represents a hitchhiking complex. Bars represent mean ± s.e.m. *P* values were calculated by unpaired two-tailed *t* test with Welch’s correction. **d**, Representative kymograph for the ΔIWS1 condition showing frequent histone rewrapping. Data are presented in the same way as in (**b**).

The other elongation factor whose omission caused a strong effect on the histone fate is IWS1^25,26^. However, unlike ΔSPT6, ΔIWS1 did not induce histone eviction but markedly increased the rewrapping frequency—either at the initial EC-nucleosome encounter or during hitchhiking (**Fig. 2a, 2d, and S4c**). This indicates the presence of IWS1 within the complete EC, counterintuitively, inhibits nucleosome reassembly, arguing against a canonical histone chaperoning activity for IWS1. Furthermore, omission of both SPT6 and IWS1 restored the histone fate distribution close to the All-Factor condition (ΔSPT6&ΔIWS1 in **Fig. 2a, S4a, and S4c**), underscoring the mutual regulation of this pair of factors in chromatin maintenance^27,28^.

Next, we sought to use an orthogonal approach to corroborate the above single-molecule results. We assembled ECs on an FAM-labeled nucleic acid scaffold, which was designed to contain minimal flanking DNA. We then incubated the scaffold with LD655-labeled histone octamers and employed native electrophoretic mobility shift assay (EMSA) to evaluate the level of histone-EC association, which could only occur via protein-protein interactions (**Fig. S5a**). Control experiments showed that the overlapping FAM and LD655 signals relied on the existence of histone octamers and that the EC with All-Factor was much more proficient in recruiting histones than Pol II alone (**Fig. S5b**). Among the elongation factors, SPT6 and IWS1 exerted the most significant effects on histone retention in this bulk assay (**Fig. S5c**): ΔSPT6 and ΔIWS1 reduced and increased the level of histone association with EC, respectively, whereas ΔSPT6&ΔIWS1 brought the level close to the All-Factor condition. The same trend was observed whether H2B or H4 was labeled in this assay (**Fig. S5d**). Together, these results show that SPT6 mediates histone association with the EC while IWS1 prevents the histones from being rewrapped onto the DNA, both promoting the formation of the hitchhiking complex.

### FACT and NAP1L1 cooperate to facilitate histone rewrapping

Since neither SPT6 nor IWS1 seems to mediate histone rewrapping after EC passes a nucleosome—in fact, we show here that IWS1 plays an inhibitory role—we turned our attention to the other factors in our system, namely, the two histone chaperones FACT and NAP1L1^29–31^. We first performed single-molecule experiments under the ΔFACT condition. We found ΔFACT only modestly increased the blocking ratio and the pausing time of EC at the nucleosome (**Fig. S2a and S2c**), but caused a significant reduction in the frequency of histone rewrapping and a corresponding elevation in histone hitchhiking frequency compared to the All-Factor condition (**Fig. 3a and S6a**). A similar phenotype was observed when NAP1L1 was omitted. Omission of both chaperones (ΔFACT&ΔNAP1L1) completely abolished the rewrapping events; as a result, essentially all histone H4 entered the hitchhiking mode (**Fig. 3a and S6a**). Notably, ΔFACT, ΔNAP1L1, and ΔIWS1 all increased the frequency of rewrapping or hitchhiking at the expense of the other but did not noticeably affect the level of H4 eviction, unlike ΔSPT6 which induced significant H4 loss (**Fig. 2a and 3a**).

**Figure 3.**
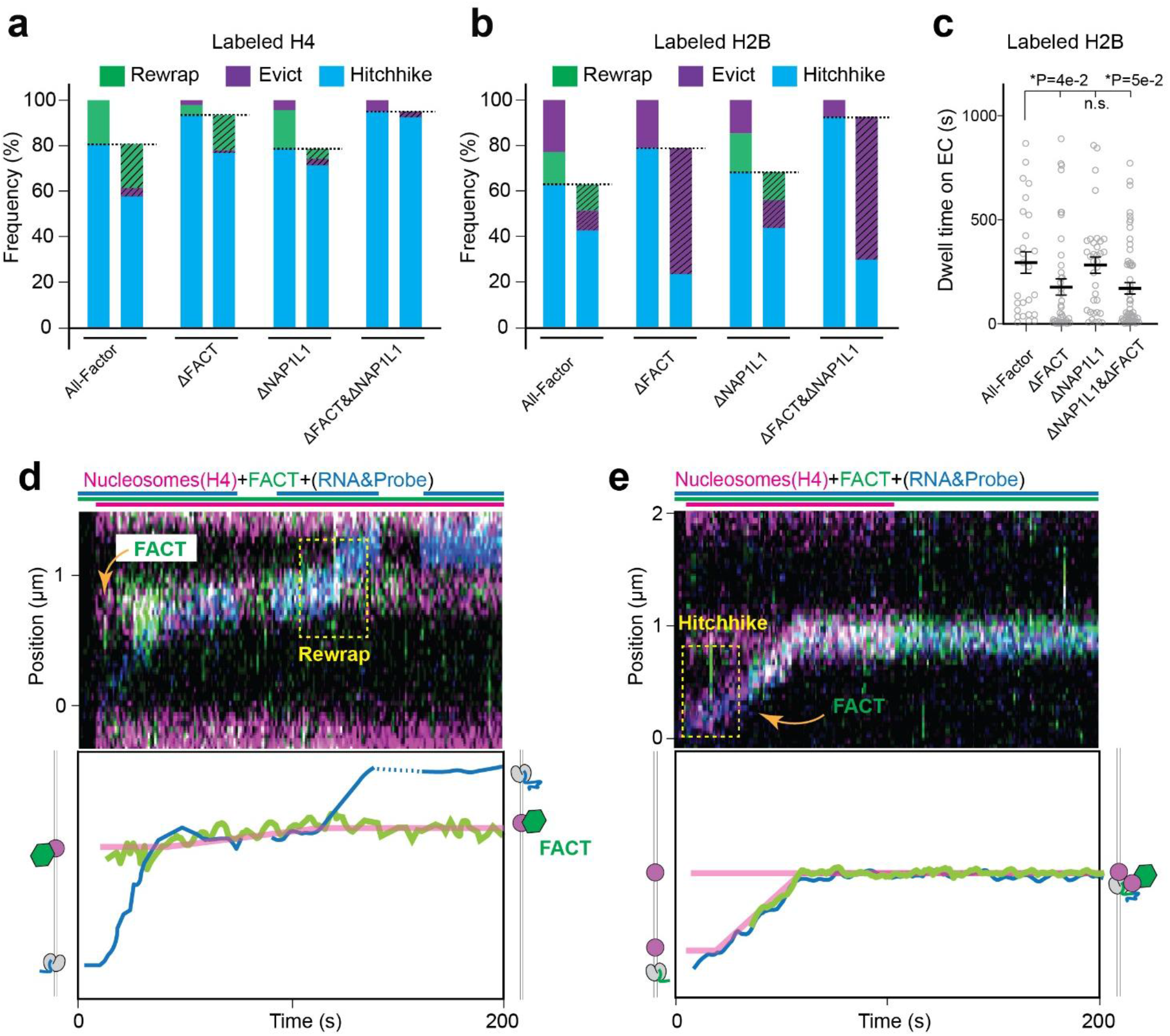
FACT and NAP1L1 promote histone rewrapping before and after hitchhiking. **a**, Frequency of different histone H4 fates after an EC passed a nucleosome under the indicated factor conditions. For each condition, both the initial histone fate upon EC-nucleosome encounter (left) and the final histone fate for a hitchhiking complex at the end of imaging (right) are shown. **b**, Frequency of different histone H2B fates under the indicated factor conditions. Data are presented in the same way as in (**a**). **c**, Residence time of H2B on EC during hitchhiking under the indicated factor conditions. Each point represents a hitchhiking complex. Bars represent mean ± s.e.m. *P* values were calculated by unpaired two-tailed *t* test with Welch’s correction. **d**, Representative kymograph showing FACT (green) prebound to a nucleosome (magenta) mediating a rewrapping event after the passage of EC (blue). **e**, Representative kymograph showing FACT being recruited to a hitchhiking complex and co-migrating with it. NAP1L1 was not included in these experiments. Bars above the kymographs indicate the time periods when specific lasers were switched on or off. Plots below the kymographs show the extracted EC, FACT, and histone positions to assist with visualization.

Thus far we have been using H4 as a proxy for the histone core. To more comprehensively characterize the histone fates, we also conducted single-molecule experiments using LD655-labeled histone H2B. Under the All-Factor condition, H2B also adopted heterogeneous behaviors (hitchhiking/eviction/rewrapping/blocking) upon EC-nucleosome encounter but displayed a higher eviction frequency than H4 (**Fig. 3a, 3b, S2c, and S6c-e**), consistent with the notion that the H2A-H2B dimer is more labile than the (H3-H4)_2 t_etramer during transcription^32^. We found that ΔFACT largely eliminated H2B redeposition (**Fig. 3b and S6a**) and caused a significant reduction in the H2B residence time on EC during hitchhiking, resulting in more eviction (**Fig. 3c and S6b**). In contrast, ΔNAP1L1 only had minor effects on the H2B fate (**Fig. 3b,c and S6a,b**). Overall, these results suggest that FACT and NAP1L1 additively facilitate redeposition of the (H3-H4)_2 t_etramer during transcription, but play differential roles in retaining the H2A-H2B dimer on EC.

Given FACT’s prominent role, we additionally labeled this complex with a Cy3 dye to directly visualize its behavior during chromatin transcription. We did not include NAP1L1 in these experiments so that FACT was the only chaperone that could mediate histone rewrapping. We found that FACT readily binds nucleosomes as expected^33^ (**Fig. S7a**), but does not bind EC on its own (0/30 events) (**Fig. S7b**). However, once an EC encountered a nucleosome and entered the hitchhiking mode, FACT was observed to frequently associate and co-travel with the hitchhiking complex (18/20 events), either prebound to the nucleosome (**Fig. 3d**) or recruited de novo to the histone-bound EC (**Fig. 3e**), potentially mediating chromatin restoration within gene bodies^34^. Indeed, we observed all of the nucleosome rewrapping events in these experiments exhibited FACT signal colocalized with the EC (6/6 events), supporting the importance of FACT in the histone hitchhiking-to-rewrapping transition.

### SPT6, IWS1, and FACT regulate chromatin organization in gene bodies

Having defined the distinct roles for SPT6, IWS1, and FACT in histone retention and rewrapping in our in vitro single-molecule assays, we next asked whether the corresponding effects could be detected in cells. We first generated human DLD-1 cells carrying acute-depletion tags on endogenous SPT6 and IWS1, enabling rapid and selective depletion of either factor within the same cellular background. Efficient depletion was confirmed by immunoblotting (**Fig. 4a**). We then performed micrococcal nuclease sequencing (MNase-seq) to map the chromatin landscape across promoter-proximal regions and early gene bodies, focusing primarily on the +1 nucleosome, the first major barrier that Pol II encounters upon entry into productive elongation in higher eukaryotes^35,36^. Such focus was intended to minimize secondary effects arising from downstream elongation defects caused by factor depletion.

**Figure 4.**
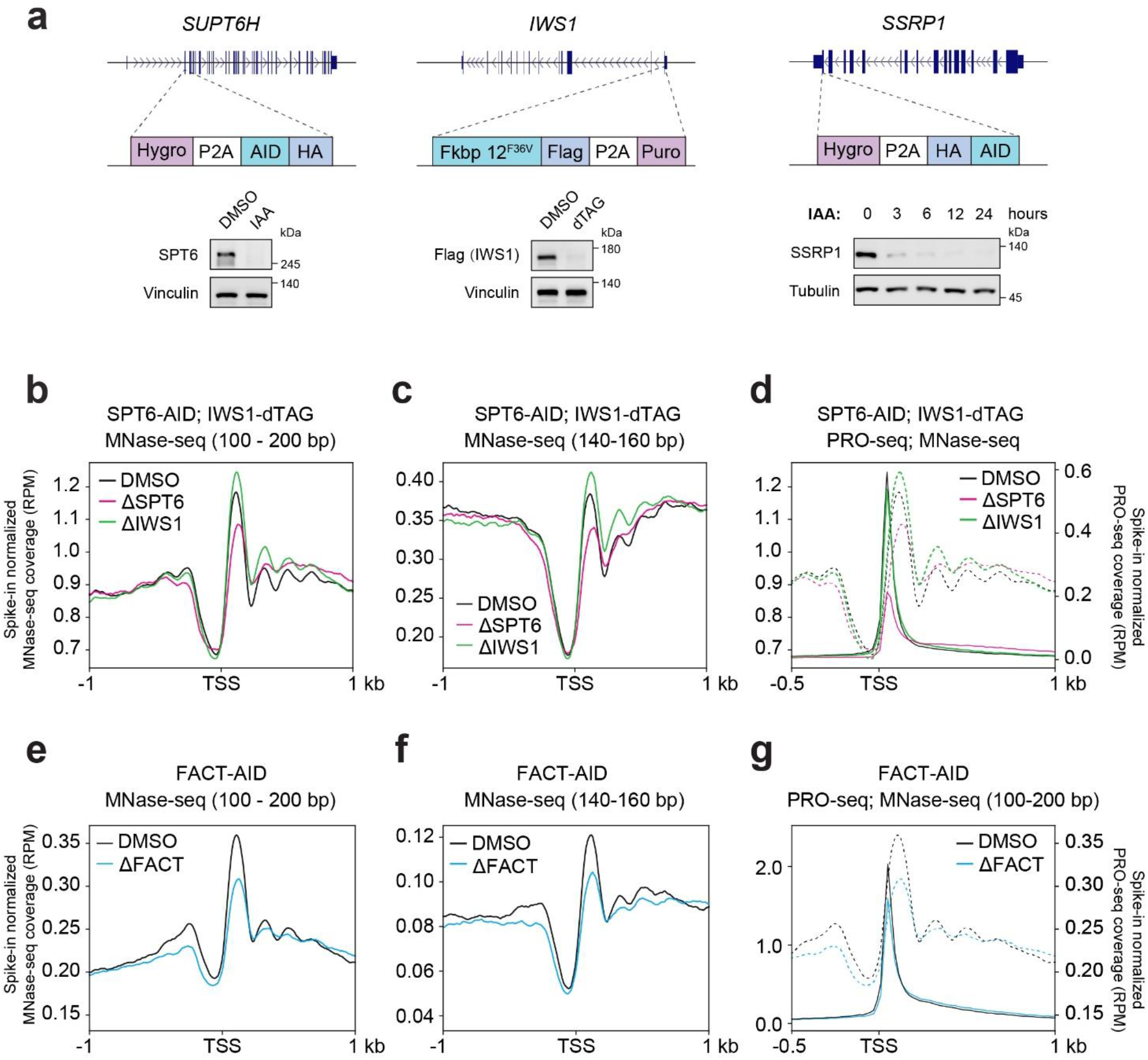
Acute depletion of SPT6, IWS1, or FACT differentially alters promoter-proximal chromatin organization in cells. **a**, Schematics showing the generation of SPT6-AID, IWS1-dTAG, and SSRP1-AID (hereafter FACT-AID) human DLD-1 cell lines by endogenous degron tagging. Western blots of whole-cell extracts from SPT6-AID cells treated with DMSO or 500 µM indole-3-acetic acid (IAA) for 3 h, IWS1-dTAG cells treated with DMSO or 200 nM dTAG-13 (hereafter dTAG) for 3 h, and FACT-AID cells treated with 1 mM IAA for the indicated durations are shown. Vinculin is a loading control for the SPT6 and IWS1 blots, and tubulin for the SSRP1 blot. A 6-h IAA treatment was used for subsequent experiments in FACT-AID cells. **b-c**, Metaplots showing the average spike-in–normalized MNase-seq coverage centered at transcription start sites (TSSs) in DMSO-treated control cells, SPT6-AID cells treated with IAA (ΔSPT6), and IWS1-dTAG cells treated with dTAG (ΔIWS1). The population of 100–200-bp MNase-resistant DNA fragments (**b**) and the 140–160-bp subset enriched for canonical nucleosome-sized particles (**c**) were analyzed. **d**, Metaplots showing the average spike-in–normalized PRO-seq coverage for sense transcription centered at TSSs under DMSO, ΔSPT6, and ΔIWS1 conditions (solid lines, right y axis) overlaid with the MNase-seq coverage for 100–200-bp fragments shown in (**b**) (dashed lines, left y axis). **e-f**, Metaplots showing the average spike-in–normalized MNase-seq coverage centered at TSSs for the 100–200-bp (**e**) and 140–160-bp (**f**) fragments in FACT-AID cells treated with DMSO or IAA (ΔFACT). **g**, Metaplots showing the average spike-in–normalized PRO-seq coverage for sense transcription centered at TSSs under DMSO and ΔFACT conditions (solid lines, right y axis) overlaid with the MNase-seq coverage for 100–200-bp fragments shown in (**e**) (dashed lines, left y axis).

We first analyzed 100–200-bp MNase-protected fragments, a population encompassing canonical nucleosomes, subnucleosomal particles, and larger chromatin-associated assemblies. Control cells displayed the expected nucleosome-depleted region at the TSS, a prominent +1 nucleosome peak, and phased downstream nucleosomes (DMSO; **Fig. 4b and S8a**). In comparison, SPT6-depleted cells (ΔSPT6) exhibited much reduced protection at the +1 nucleosome and disrupted downstream nucleosome phasing, consistent with previous work^37^. On the contrary, ΔIWS1 increased the signal at the +1 position (**Fig. 4b and S8b,c**). To narrow down the specific chromatin species underlying these opposing effects, we stratified the fragments by size and focused on the 140–160-bp subpopulation, which is enriched for canonical nucleosome-sized particles. We found this subpopulation was again reduced at the +1 position for ΔSPT6 but increased for ΔIWS1 (**Fig. 4c**). By contrast, fragments shorter than 140 bp increased following depletion of either factor (**Fig. S8d**). Thus, SPT6 loss in the cell shifted the +1 nucleosome population from canonical nucleosome-sized particles toward shorter protected intermediates, consistent with defective histone retention, while IWS1 loss increased both canonical and shorter protected particles, consistent with enhanced histone–DNA reassociation that populates both fully and partially wrapped states.

Because the +1 nucleosome contributes to the stabilization of promoter-proximally paused Pol II^38^, we next inquired whether the changes in nucleosome organization caused by ΔSPT6 and ΔIWS1 are accompanied by altered Pol II pausing. We performed precise run-on sequencing (PRO-seq) experiments and observed the expected sharp accumulation of transcriptionally engaged Pol II immediately downstream of the TSS and upstream of the +1 nucleosome in control cells (**Fig. 4d and S8e**). ΔSPT6 dramatically reduced this Pol II peak, consistent with the loss of canonical +1 nucleosome occupancy. This promoter-proximal redistribution agrees with previous acute-depletion studies showing reduced Pol II occupancy at pause sites or its displacement toward the +1 nucleosome region^37,39,40^. ΔIWS1 had little effect on paused Pol II despite increasing protection of nucleosomes and subnucleosomal particles (**Fig. 4d and S8f**), perhaps because Pol II pausing at the +1 nucleosome is already maximized in the wildtype condition. This limited promoter-proximal effect is compatible with recent mNET-seq data showing that acute IWS1 depletion mainly causes reduced elongation velocity and Pol II accumulation within gene bodies^41^.

We then used a similar strategy to acutely deplete FACT in the cell (**Fig. 4a**). MNase-seq data showed that ΔFACT caused a pronounced reduction in the 100–200-bp and 140–160-bp fragment protection at the +1 nucleosome and more modest changes at downstream nucleosomes (**Fig. 4e,f and S8g,h**). PRO-seq data further showed a reduction in the promoter-proximally paused Pol II peak following FACT depletion, consistent with previous observations^42^ (**Fig. 4g and S8i**). These findings support a direct role of FACT in local nucleosome restoration following Pol II passage. Together, these cellular data are consistent with the in vitro single-molecule results: SPT6 retains displaced histones on EC, FACT promotes their rewrapping onto DNA, and IWS1 restrains spurious histone rebinding. The balance among these activities preserves the promoter-proximal nucleosome landscape and the associated paused Pol II state.

### The hitchhiking EC unloads histones before traversing the next nucleosome

Our experiments identified the hitchhiking complex to be a prevalent and stable form of EC during chromatin transcription. This finding begs a new question: can an EC keep carrying additional histones as it moves along nucleosome arrays? The answer lies in the single-molecule trajectories where EC encountered consecutive nucleosomes. We found that once a hitchhiking EC met a downstream nucleosome, it was predominantly blocked (∼80%) (**Fig. 5a and 5b**). This is in stark contrast to the efficient passage of an unoccupied EC across the first nucleosome. Furthermore, we found that the hitchhiking ECs translocated on DNA at similar speeds compared to free ECs before encountering the next nucleosome (**Fig. 5c**). These results suggest that the histones bound to EC do not substantially impair Pol II elongation on DNA but rather compromise its ability to unravel and traverse the next nucleosome.

**Figure 5.**
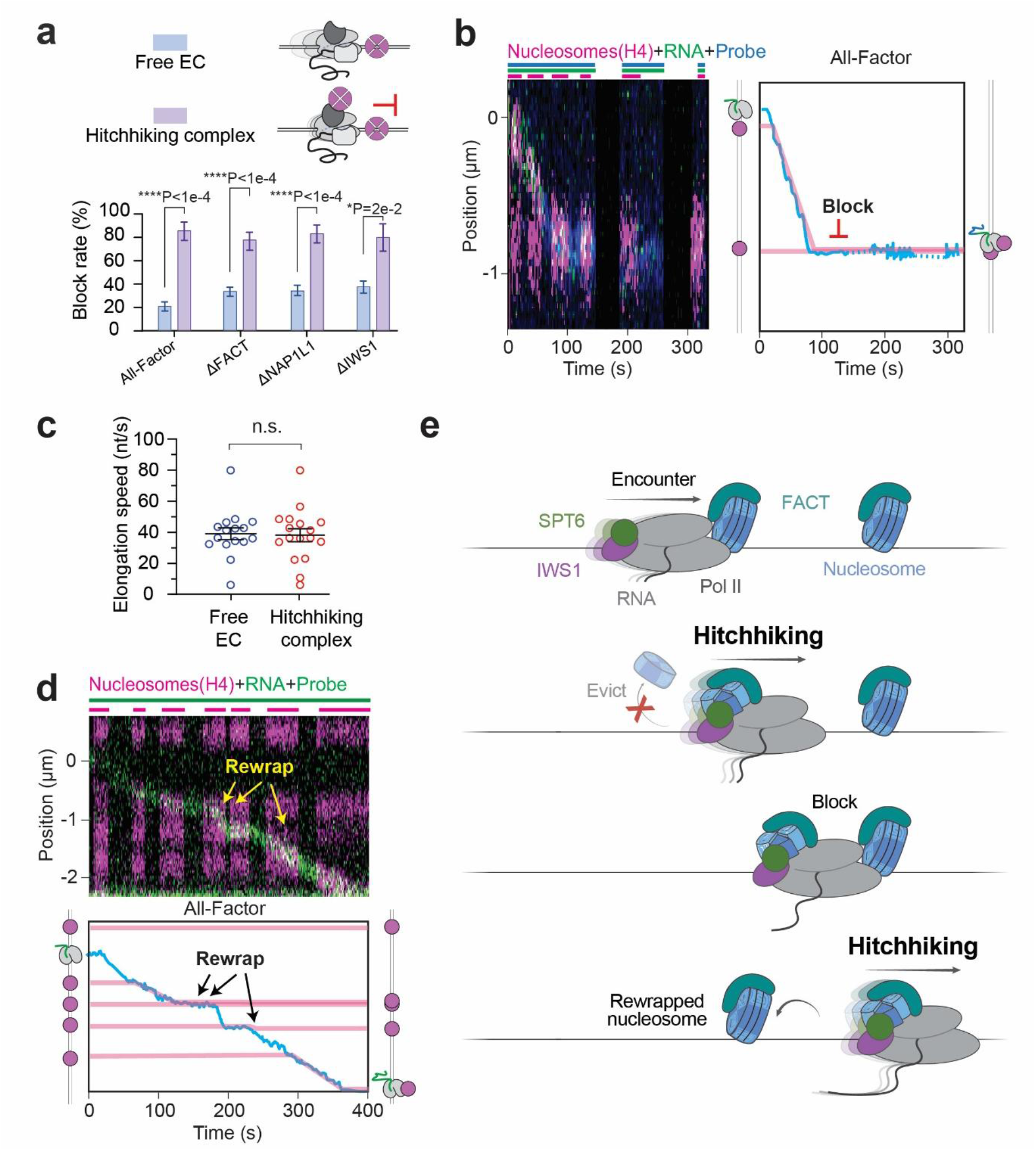
The next nucleosome presents a strong barrier to a hitchhiking complex. **a,** Fraction of ECs free of any associated histones (blue) or in the form of a hitchhiking complex (purple) that were blocked by a downstream nucleosome under the indicated factor conditions. Bars represent mean ± s.e.m. *P* values were calculated using two-sided Fisher’s exact tests. **b,** (Left) Representative kymograph showing a hitchhiking complex being blocked by a downstream nucleosome. (Right) EC trajectory (cyan) and histone positions (magenta) are extracted from the kymograph to assist with visualization. **c**, Average elongation speed for free ECs (blue) and hitchhiking complexes (red) under the All-Factor condition. Each point represents an EC. Bars represent mean ± s.e.m. *P* values were calculated by unpaired two-tailed *t* test with Welch’s correction. **d**, (Left) Representative kymograph showing an EC traversing multiple nucleosomes, each time accompanied by unloading and rewrapping of its associated histones. (Right) Extracted EC trajectory (cyan) and histone positions (magenta). **e**, Working model for transcription-coupled chromatin maintenance.

How can ECs then traverse multiple nucleosomes as they must do within active genes? We did record single-molecule trajectories where an EC managed to progress through successive nucleosomes (**Fig. 5d and S9a,b**). In the vast majority of cases (51/59 events), the EC was observed to unload its bound histones—mostly via redeposition onto DNA (45/51)—either before or at the next nucleosome encounter. This finding suggests that the conversion of a hitchhiking EC back to a free EC is essential for its successful traversal through the next nucleosome. It further predicts that EC should have a lower probability to pass multiple nucleosomes under conditions that hinder histone unloading. Indeed, the ΔFACT&ΔNAP1L1 condition, which abolished histone rewrapping (**Fig. 3a**), resulted in severely impaired second-nucleosome passage compared to the All-Factor condition (**Fig. S9c**).

## Discussion

Previous studies suggested nucleosome transfer from downstream to upstream DNA upon Pol II traversal occurs either via template looping or mediated by FACT^43,44^, but the detailed mechanism remains unresolved. By directly observing the dynamic behavior of actively transcribing ECs and the nucleosomes on their path, our study uncovers a prevalent intermediate which we termed the hitchhiking complex (i.e., a translocating EC carrying the dislodged histones.) This finding has major implications for the mechanism of chromatin transcription (**Fig. 5e**). First, EC is not obligated to rewrap the displaced histones in situ but has the tendency to carry them and continue translocating downstream. As such, the EC-histone complex is not a transient but a stable assembly, thereby ensuring robust histone retention. Second, the timing of nucleosome reassembly is influenced by several intertwining factors. As such, the EC is not obligated to rewrap the dislodged histones right away but has multiple opportunities to reassemble the nucleosome. Moreover, the positioning of the next nucleosome provides an additional checkpoint to prevent causing long-range chromatin disruption.

Our work identified several key factors that control the stability and resolution of the hitchhiking complex. We found that SPT6 contributes significantly to the retention of histones, likely mediated by its acidic and intrinsically disordered N-terminal region as suggested by recent work in yeast^45^. We envision that other acidic IDR-containing elongation factors, such as DSIF and PAF1C, may also be involved in carrying the basic histone core^46^. Notably, in the companion manuscript, we reported that the phosphorylated C-terminal domain of Pol II alone is able to carry histones^47^, indicating multiple routes towards the formation of the hitchhiking complex. Moreover, we found histone chaperones FACT and NAP1L1 counteract IWS1 and promote nucleosome reassembly, consistent with their expected roles in chromatin transcription^48,49^. In addition, our data show that FACT can be part of the hitchhiking complex and contribute to retention of the H2A-H2B dimer, although whether the EC carries one or two copies of H2A-H2B during hitchhiking requires further investigation. It will be interesting to examine how histone variants and active histone posttranslational modifications modulate the kinetic competition between histone hitchhiking and redeposition. In the companion manuscript, we found similar hitchhiking propensity using the H3.3 variant^47^, indicating the universality of this mechanism.

Our data show that the hitchhiking complex translocates on DNA at a similar velocity as a free EC but tends to be blocked by the next nucleosome. Such limited histone-carrying capacity suggests that nucleosome spacing within gene bodies (typically 30-50 bp) may constrain how far the displaced histones can be moved downstream. Previous models for transcription-coupled histone transfer based on structural data necessitate an upstream shift of the reassembled nucleosome^13,16^. Local hitchhiking within closely spaced nucleosome arrays could partially offset this shift and restore the original nucleosome positions, which may then be further refined by histone-sliding chromatin remodelers^50^. While our model depicts a faithful transcription-coupled nucleosome preservation mechanism, there is evidence that high transcriptional activities can deplete nucleosomes across gene bodies^51^. It can be envisioned that a train of ECs initiating from a strong promoter harbors a larger hitchhiking capacity and causes longer-range nucleosome repositioning compared to a single EC. This conjecture is supported by a recent study employing single-molecule chromatin fiber sequencing to probe chromatin architecture in the cell^52^.

In conclusion, by directly visualizing transcription elongation on chromatinized DNA, this study provides a framework for understanding how histones and their associated epigenetic information are retained, restored, and redistributed during Pol II transcription in higher eukaryotic cells.

## Acknowledgments

We thank members of the Liu Laboratory for discussion and G.N.L. Chua for help with the manuscript preparation. S.L. is supported by the Alfred P. Sloan Foundation, the Mathers Foundation, the Jensam Foundation, and the Marlene Hess Center for Research on Women’s Health and Biomedicine. Y.X. is supported by grants from the National Key R&D Program of China (2021YFA1300100), the National Natural Science Foundation of China (32271242), the XPLORER Prize, and the New Cornerstone Investigator Program.

## Author contributions

Y.W., X.C., Y.L., F.X.C., Y.X. and S.L. designed the experiments. Y.W. performed the single-molecule experiments and conducted data analysis with assistance from J.W.W.. X.C. prepared the protein reagents. Y.L. performed the cellular experiments and analyzed the data with assistance from A.S.. S.L., Y.W., F.X.C. and Y.X. wrote the manuscript with inputs from all authors.

## Competing interests

The authors declare no competing interests.

## Data availability

All data are available in the main text or supplementary materials.

## Methods

### Protein preparation

Pol II was isolated from *S. scrofa* thymus and *H. sapiens* TFIIS, P-TEFb, PAF1C, ELOF1, SPT6, DSIF, RTF1, and IWS1 were prepared as previously described^21^. Purified DSIF and Pol II were pre-phosphorylated by incubation with concentrated P-TEFb at approximate final concentrations of 19 μM DSIF or 3 uM Pol II and 2 μM P-TEFb. The phosphorylation reaction contained 2 mM ATP, 5 mM MgCl₂ and 5 mM freshly prepared DTT, and was incubated at 30 °C for 1 h, then aliquoted and stored at −80 °C.

### FACT

The two full-length ORFs of human FACT were cloned into a modified pCAG vector, and SUPT16H was tagged with an N-terminal protein A-SUMO tag or an N-terminal protein A-SUMO-S6 tag (S6: GDSLSWLLRLLN)^53^. The plasmid was transfected into Expi293F cells for overexpression. After cultured at 37 °C for 72 h, cells were harvested and lysed in Lysis buffer containing 30 mM HEPES pH 8.0, 300 mM NaCl, 0.25% CHAPS, 5 mM ATP, 5 mM MgCl_2,_ 0.2 mM EDTA, 2 mM DTT, 10% glycerol (v/v), 1 mM PMSF, 1 μg/mL Aprotinin, 1 μg/mL Pepstatin and 1 μg/mL Leupeptin at 4 °C for 30 min. The lysate was clarified by centrifugation at 15,000 rpm for 40 min at 4 °C. The supernatant was incubated with IgG resin for 1.5 h followed by on-column digestion by ULP1 protease at 4°C for 1 h. The immobilized proteins were eluted and further purified by a mono Q 5/50 GL column (Cytiva) pre-equilibrated in buffer A (30 mM HEPES pH 8.0, 30 mM NaCl, 2 mM DTT, 2 mM MgCl_2 a_nd 5% (v/v) glycerol). The target protein was eluted with a gradient from 0% to 100% buffer B (same as buffer A except with 1 M NaCl) over 30 column volumes. The peak fractions were pooled, aliquoted, flash-frozen in liquid nitrogen, and stored at −80 °C.

To site-specifically label the FACT complex, the S6-FACT, Sfp synthase and Cy3-CoA dye (SiChem) were incubated at a 1:1.33:11 molar ratio for 3 h at room temperature and overnight at 4 °C in the presence of 10 mM MgCl_2_. Excess dye was removed by Micro Bio-Spin Columns P30 (BIO-RAD) with a buffer containing 300 mM NaCl, Tris-HCl 20 mM pH 8.0, 5 mM MgCl_2,_ 1 mM DTT. The labeled protein was evaluated via SDS-PAGE gel electrophoresis. The gel was scanned by Typhoon FLA 7000 (GE Healthcare). The labeling efficiency was estimated to be 70%. Final protein samples were aliquoted, flash frozen, and stored at −80 °C.

### hNAP1L1, yNap1

Recombinant *Saccharomyces cerevisiae* Nap1 (yNAP1) was expressed and purified as previously described^54^. For human NAP1L1 (hNAP1L1), the full-length ORF of hNAP1L1 was cloned into a modified pGEX-6p-1 vector containing an N-terminal GST-3C tag and expressed in *E. coli* BL21 (DE3) cells. Cells grew at 37 °C until reaching an optical density at 600 nm (OD_600)_ of around 0.6. The temperature decreased to 16 °C and protein expression was induced by adding 0.1 mM Isopropyl β-D-1-thiogalactopyranoside (IPTG). Cells were grown for an additional 16 h at 16 °C and were harvested and re-suspended in a buffer containing 20 mM HEPES 7.6, 500 mM NaCl, 0.05% CHAPS and 1 mM DTT. Cells were lysed by high pressure at 4 °C for 20 min and the lysate was cleared by centrifugation at 15,000 rpm for 40 min at 4 °C. The supernatant was incubated with GST resin for 1 h followed by on-column digestion by HRV-3C protease at 4°C overnight. The immobilized protein was eluted and further purified by a mono Q 5/50 GL column (Cytiva) pre-equilibrated in buffer A (20 mM HEPES pH 7.6 and 1 mM DTT). The target protein was eluted with a gradient from 0% to 100% buffer B (same as buffer A except with 1 M NaCl) over 30 column volumes. The peak fractions were pooled, aliquoted, flash-frozen in liquid nitrogen, and stored at −80 °C.

Histone proteins

Recombinant human core histones were prepared and fluorescently labelled with LD655 essentially as described previously^55^. Briefly, individual core histones were expressed in E. coli BL21(DE3) pLysS cells, isolated from inclusion bodies and purified under denaturing conditions by sequential Q and SP ion-exchange chromatography. Site-specific labelling was performed under denaturing conditions by reacting H4 L50C or H2B K120C with LD655-maleimide. Histone octamers were assembled by combining equimolar amounts of the four core histones, including either LD655-labelled H4 L50C or LD655-labelled H2B K120C together with the corresponding unlabeled histones, followed by refolding and purification by size-exclusion chromatography as described previously. The labeling efficiency of H4 and H2B were estimated to be 71% and 82%, respectively. Final protein samples were aliquoted, flash frozen, and stored at −80 °C.

### DNA substrate preparation

DNA substrates were generated essentially as described previously^21^. In brief, 8-kb plasmid templates containing a unique BbsI site and a downstream repeat cassette were linearized, modified to generate extended single-stranded overhangs, and terminally biotinylated by Klenow-mediated fill-in with biotinylated nucleotides. Template- or non-template-strand oligonucleotides were then ligated to the biotinylated DNA to generate TS-DNA and NTS-DNA handles for elongation complex assembly. The final products were purified by size-selective magnetic beads, aliquoted and stored at −80 °C.

### Single-molecule experiments

Elongation complexes were assembled same as described previously^21^. Briefly, biotinylated template-strand DNA handles were annealed with a fluorescently labelled RNA primer (Cy3 or AF488) (Integrated DNA Technologies) to generate the initial DNA–RNA scaffold. Pre-phosphorylated Pol II, was then added to the annealed scaffold to form a partial EC. The corresponding non-template-strand DNA handle was subsequently added to complete EC assembly. Annealing-generated nicks were sealed with T4 DNA ligase, and the assembled ECs were loaded directly into microfluidic chambers for single-molecule assays.

Single-molecule experiments were performed at room temperature using a LUMICKS C-Trap instrument. The microfluidic channels of a C2 flow cell were passivated by sequential incubation with 0.1% (w/v) BSA in 1× PBS and 0.5% (w/v) Pluronic F127 in 1× PBS for 10 min each. Streptavidin-coated polystyrene beads (2.14 μm in diameter; Spherotech), 16-kb DNA-ECs (100 pM in imaging buffer), imaging buffer, nucleosome-loading buffer and transcription buffer were introduced into channels 1–5, respectively. Imaging buffer contained 20 mM Tris-HCl, pH 8.0, 40 mM KCl, 2 mM MgCl_2 a_nd 500 μg ml⁻¹ BSA, and was supplemented with an oxygen-scavenging system (10 nM protocatechuate-3,4-dioxygenase and 2.5 mM protocatechuic acid), 0.3 U μl⁻¹ murine RNase inhibitor, 5 mM freshly prepared DTT and a triplet-state-quenching cocktail containing 1 mM cyclooctatetraene and 1 mM 4-nitrobenzyl alcohol. Transcription buffer was prepared in imaging buffer and contained all four ribonucleoside triphosphates (rNTPs), each at 1 mM, together with the indicated elongation factors and histone chaperones. For the all-factor condition, the final factor concentrations were as follows: PAF1C 100 nM, SPT6 120 nM, RTF1 400 nM, TFIIS 2 μM, pre-phosphorylated DSIF 200 nM, IWS1 120 nM, ELOF1 3 μM, P-TEFb 50 nM, FACT 250 nM, and NAP1L1 50 nM. The final salt and buffer composition of the transcription buffer was 16 mM Tris-HCl (pH 8.0), 11 mM HEPES (pH 8.0), 62 mM NaCl, 16 mM KCl and 2.1 mM MgCl_2_. In experiments using fluorescently labelled FACT, Cy3-FACT was used at 20 nM to minimize fluorescence background. Nucleosome-loading buffer contained 3 nM LD655-labelled histone octamers and either 4 nM yNap1 or 4 nM NAP1L1 in 1× HR buffer containing 30 mM Tris-HCl, pH 7.5, 50 mM KCl, 1 mM freshly prepared DTT and 50 μg ml⁻¹ BSA.

Single DNA tethers were formed between two streptavidin-coated beads held in optical traps; the tethers were maintained at a constant tension of 3 pN. Tethers were moved to channel 3, containing imaging buffer, for confocal scanning with either 561- or 488-nm excitation to verify the presence of an assembled EC. Tethers lacking a detectable EC signal or containing protein aggregates were discarded. For chaperone-assisted loading, nucleosome-containing DNA tethers were generated using yNap1 or hNAP1L1 as previously described^56^. Briefly, EC-containing DNA tethers were moved into channel 4 containing nucleosome-loading buffer, the chaperone and histone octamers. After incubation for 1–3 s, the tether was transferred to channel 5 containing transcription buffer. For salt-dialysis-mediated loading, the NTS-DNA (10 nM final concentration) was mixed with histone octamers (400-600 nM final concentration) and dialyzed as previously described^55^. The octamer concentration was optimized to yield a few nucleosomes per DNA tether. EC-containing nucleosome-loaded DNA tether was directly transferred to channel 5. Kymographs were acquired using Bluelake software (LUMICKS) by confocal line scanning through the centers of the two trapped beads at 0.2 s per line, with a pixel dwell time of 0.2 ms. Data acquisition was initiated in imaging-buffer channel 3 for 10 s to establish a baseline and continued throughout subsequent measurements in channels 4 and 5. Where indicated, an AF488-or Cy3-labeled DNA oligonucleotide probe (7 nM; 5′-Dye-AGTGTGTGGTCTGTGGTGTCT-3′; Integrated DNA Technologies) was included in the transcription buffer to detect nascent RNA.

### Single-molecule data analysis

Single-molecule force and fluorescence data from .h5 files generated by Bluelake were processed using the lumicks.pylake Python library, supplemented by additional Python modules integrated into a custom graphical user interface script, CTrapVis.py (http://harbor.lumicks.com/single-script/c5b103a4-0804-4b06-95d3-20a08d65768f). This script was used to export confocal scans and kymographs in TIFF format and to extract time-resolved force, distance and photon-count data. The positions of labelled RNA and nascent-RNA probes on the DNA tether over time were determined from kymographs using a custom script, kymotracker_calling_script.py (https://harbor.lumicks.com/single-script/4db9d63e-1f93-0c0-9b90-e99066469578), which implemented the greedy line-tracking algorithms provided by lumicks.pylake to define line traces. Optimal tracking of EC trajectories was achieved by setting a pixel threshold of 2 photon count for Cy3 signals and 1 photon count for AF488 signals, with a line width of 12 pixels, a window of 35 frames and a minimum line length of 20 frames. Extracted trajectories were aligned based on the starting position of the labelled RNA signal collected before transcription initiation. Distance travelled in nanometers was converted to base pairs using a scaling factor of 0.285 nm per bp, reflecting the extension of B-form DNA under 3 pN of tension. Nucleosome fates were assigned by visual inspection of multi-color kymographs, as the different outcome classes were directly distinguishable from the relative trajectories of the EC and LD655-labelled nucleosome signals. Block events were defined as encounters in which the EC became colocalized with a nucleosome and remained immobile at the nucleosome position for more than 300 s. Pairwise comparisons of single-molecule outcome frequencies were performed using two-sided Fisher’s exact tests. For each comparison, the number of events in the indicated category was compared with the number of all remaining events between two conditions.

Segmentation analysis of extracted transcription trajectories was performed essentially as described previously^21^. Briefly, EC trajectories obtained from kymographs were segmented into transcriptional intervals, and elongation speeds were calculated from the slopes of individual segments after converting distance travelled in nanometers to base pairs. For the analysis in Fig. 5c, trajectory segments were assigned to empty ECs, ECs carrying one hitchhiked nucleosome based on direct visual inspection of multicolor kymographs. These states were readily distinguished by the presence or absence of LD655-labeled nucleosome signals colocalized with the moving EC, and therefore did not require automated classification.

### Electrophoretic mobility shift assay (EMSA)

Two-color electrophoretic mobility shift assays were performed using the same template strand, non-template strand, and RNA scaffold as in a previously described protocol^57^. The template strand was 5′-FAM-labeled to visualize the Pol II elongation complex, whereas histone octamers were labeled with LD655 on either H4 or H2B to monitor octamer association with the EC. To assemble ECs, FAM-labeled template strand and RNA were first annealed at a molar ratio of 1:1 using the following annealing procedure: 55 °C for 30 s, 50 °C for 30 s, 45 °C for 30 s, 40 °C for 2 min, 37.5 °C for 2 min, 35 °C for 2 min, 30 °C for 2 min, 27 °C for 2 min, 25 °C for 10 min, followed by a hold at 4 °C. The annealed template strand–RNA hybrid was then mixed with pre-phosphorylated Pol II at a TS:Pol II molar ratio of 1:1.3 and incubated at 30 °C for 10 min in the presence of RNase inhibitor and freshly prepared DTT. Non-template strand was subsequently added to a final TS:NTS molar ratio of 1:1.5, followed by incubation at 30 °C for 10 min and cooling to 4 °C. For binding reactions, ECs were mixed with the indicated elongation factors or factor combinations and incubated for 5 min at 30 °C followed by 5 min at 4 °C. LD655-labeled histone octamer, containing either H4-LD655 or H2B-LD655, was then added, followed by an additional incubation for 5 min at 30 °C and 5 min at 4 °C. DSIF was pre-phosphorylated by P-TEFb before use. Unless otherwise indicated, final reactions contained 100 nM EC, 200 nM LD655-labeled histone octamer and 200 nM of each elongation factor. Samples were resolved on NativePAGE 3–12% Bis-Tris gels. Electrophoresis was performed in 1× Tris-glycine running buffer, pH 8.3, at 220–300 V for 2 h. Gels were scanned using a Typhoon FLA 9500 imager to detect FAM and LD655 fluorescence. The resulting LD655/FAM ratios were then normalized to the All-Factor condition for plotting.

### Cell lines

Human DLD-1 cells were cultured in McCoy’s 5A medium (BasalMedia) supplemented with 10% fetal bovine serum (Excell) and 1× penicillin–streptomycin (Gibco, Thermo Fisher Scientific) at 37 °C in a humidified incubator with 5% CO₂.

Endogenous degron knock-in and clonal isolation were performed using a CRISPR/Cas9-assisted precise integration into target chromosome (PITCh) strategy, as described previously^58–60^. Briefly, 2 μg PITCh cutting sgRNA and 4 μg donor plasmid were mixed with 2 × 10^6^ DLD-1 cells followed by electroporation using a Neon transfection system (Thermo Fisher Scientific). Following electroporation, cells were immediately transferred to 6-well plates containing prewarmed, antibiotic-free McCoy’s 5A medium and allowed to recover for 2 days. Cells were then subjected to limiting dilution and cultured for 10–14 days in McCoy’s 5A medium containing either 1 μg/mL puromycin (Meilunbio) or 400 μg/mL hygromycin B, as appropriate. Resistant colonies were transferred to 96-well plates and clonally expanded before genotyping by PCR. Degradation of the tagged proteins in homozygous knock-in clones was assessed by western blotting after treatment with 200 nM dTAG-13 for 3 h, 500 μM IAA for 3 h, or 1 mM IAA for 6 h, as appropriate; DMSO-treated cells served as controls.

### Micrococcal nuclease sequencing (MNase-seq)

MNase-seq was performed with modifications to published protocols^61^. For each biological replicate, 1 × 10^6^ experimental cells were mixed at harvest with 2 × 10^5^ mE14 mouse embryonic stem cells as spike-in controls. The pooled cells were washed with ice-cold phosphate-buffered saline and incubated in HM2 buffer containing 10 mM HEPES (pH 7.4), 2 mM MgCl₂, 0.5% NP-40, and 1× protease inhibitor cocktail (Roche) for 5 min on ice. Nuclei were then isolated, and chromatin was digested with 0.1 μL micrococcal nuclease (New England Biolabs) in the presence of 2 mM CaCl₂ for 10 min at 37 °C. The reaction was terminated by adding an equal volume of 2× stop solution containing 20 mM Tris–HCl (pH 8.0), 300 mM NaCl, 20 mM EDTA, and 4 mM EGTA. Samples were then incubated with 3 μL of RNase A (Thermo Fisher Scientific) for 30 min at 37 °C, followed by treatment with 20 μg/mL proteinase K (Beyotime) for 30 min at 55 °C. DNA was purified by phenol–chloroform–isoamyl alcohol extraction followed by ethanol precipitation.

Sequencing libraries were prepared using the VAHTS Universal Plus DNA Library Prep Kit (Vazyme) according to the manufacturer’s instructions. Library quality was assessed using an Agilent 2100 Bioanalyzer. Libraries were sequenced on a DNBSEQ-T7 platform to generate 150-bp paired-end reads.

### Precision run-on sequencing (PRO-seq)

PRO-seq was performed according to established protocols^62,63^, with modifications. Briefly, DLD-1 cells were permeabilized, resuspended in freezing buffer, flash-frozen in liquid nitrogen, and stored at −80 °C. *Drosophila* nuclei were added to each reaction at 20% of the DLD-1 input as a spike-in control. Run-on reactions were performed with biotin-11-NTPs for 5 min at 37 °C. Nascent RNA was extracted with TRIzol LS Reagent (Thermo Fisher Scientific), fragmented with 0.25 M NaOH, purified using the HiPure RNA Pure Micro Kit (Magen), and ligated to a 3′ RNA adapter. Biotinylated RNA was enriched using streptavidin magnetic beads (New England Biolabs) and sequentially subjected to on-bead 5′-hydroxyl repair with T4 polynucleotide kinase (NEB), 5′-end decapping with RppH (NEB), and 5′ RNA adapter ligation. RNA was recovered with TRIzol (Thermo Fisher Scientific), reverse-transcribed using Maxima H Minus Reverse Transcriptase (Thermo Fisher Scientific), amplified by PCR, and purified using DNA Clean Beads (Vazyme). Library quality was assessed using an Agilent 2100 Bioanalyzer, and libraries were sequenced on an Illumina NovaSeq 6000 platform in paired-end mode.

### Genomic data analysis

#### Identification of transcription start sites (TSSs)

Transcription start sites were defined using published PRO-cap data generated in DLD-1 cells, following our previously established approach^64^. RefSeq gene annotations were obtained from the UCSC Genome Browser. For each protein-coding transcript, PRO-cap signal was examined from 10 bp upstream to 300 bp downstream of the annotated TSS, and the position with the highest PRO-cap signal was defined as the refined TSS. For genes with multiple transcripts, the transcript with the highest PRO-cap signal was selected as the representative transcript.

#### MNase-seq analysis

Paired-end MNase-seq reads were trimmed with Trim Galore v0.6.6 (https://www.bioinformatics.babraham.ac.uk/projects/trim_galore/) and aligned to the human hg38 and mouse mm10 genomes using Bowtie2 v2.5.4^65^ with the --very-sensitive-local, --no-discordant, --no-mixed, and --dovetail options. PCR duplicates were removed using Picard MarkDuplicates v2.23.3 (https://broadinstitute.github.io/picard/), and properly paired alignments with MAPQ ≥ 30 were retained using SAMtools v1.21^66^. Mouse-aligned reads were used for spike-in normalization, with a sample-specific scaling factor calculated as 1e6/mm10_count. Spike-in-normalized hg38 coverage tracks were generated using deepTools bamCoverage v3.5.5^67^, with ENCODE blacklist regions excluded^68^.

To examine MNase protection profiles across fragment sizes, deduplicated hg38 alignments were separated into the indicated fragment-length groups, including 100–200 bp, 140–160 bp (canonical nucleosome-sized fragments), and 100–140 bp (shorter fragments potentially derived from subnucleosomal or other noncanonical chromatin-associated particles). Spike-in-normalized coverage tracks were generated for each fragment group using deepTools bamCoverage with centered fragments. TSS-centered heatmaps and average profiles were generated using deepTools computeMatrix, plotHeatmap, and plotProfile.

V-plots were generated from 40–250-bp fragments by plotting fragment length against midpoint position relative to TSSs, with minus-strand genes reoriented to maintain a consistent transcriptional direction. Signals were normalized to the genome-wide fragment length distribution^69^.

#### PRO-seq analysis

Raw PRO-seq reads were processed using Trim Galore (v0.6.6), and reads shorter than 15 bp after trimming were discarded. Ribosomal RNA reads were removed by alignment to the rRNA reference using Bowtie2 v2.5.4^65^, and the remaining reads were aligned to the human hg38 and fly dm6 genomes using Bowtie2 with the --very-sensitive-local option. PCR duplicates were removed using UMI-tools v1.1.6^70^, and low-quality alignments were discarded. Fly-aligned reads were used for spike-in normalization, with a sample-specific scaling factor calculated as 1e6/dm6_count. Single-base-pair-resolution, spike-in-normalized, strand-specific coverage tracks were generated using deepTools bamCoverage v3.5.5^67^ with --Offset 1 and the corresponding strand-specific SAM flags --samFlagInclude 83 or --samFlagInclude 99. TSS-centered heatmaps and average profiles were generated using deepTools computeMatrix, plotHeatmap, and plotProfile.

## Notes

### Competing Interest Statement

The authors have declared no competing interest.

