## Supplemental Material for "A histone hitchhiking mechanism for chromatin maintenance during mammalian transcription elongation"

**Supplementary Figures**

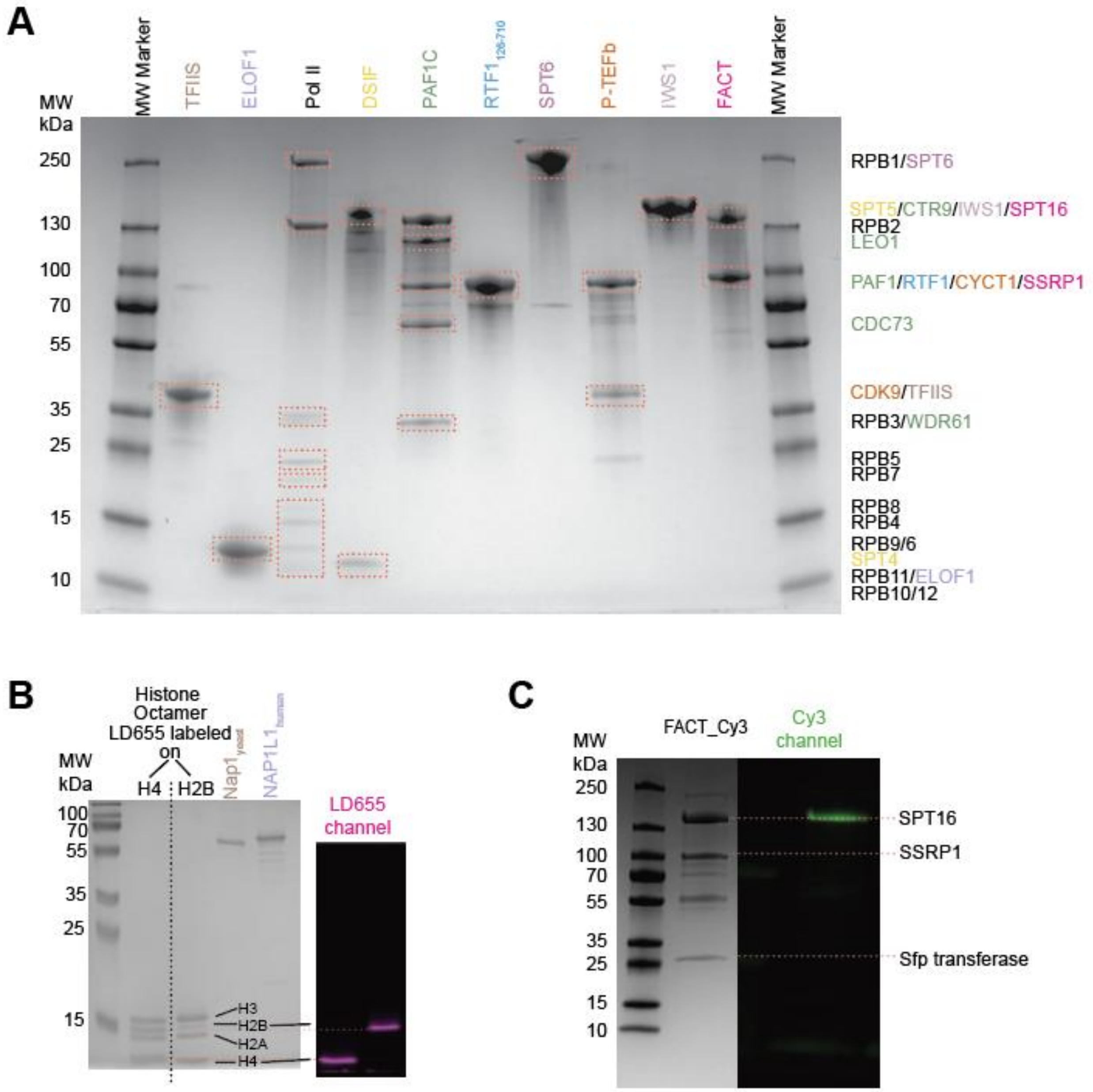


**Figure S1. Quality evaluation for protein purification and fluorescent labeling.**
(**A**) Coomassie-stained SDS-PAGE gel showing purified mammalian Pol II, elongation factors, and histone chaperones used in this study. Protein subunit identities based on their molecular weights are annotated on the right. (**B**) (Left) Coomassie-stained SDS-PAGE gel showing purified histone octamers, yeast Nap1, and human NAP1L1. Histone octamers were fluorescently labeled with LD655 on either H4 or H2B as indicated. (Right) Corresponding fluorescence scanning confirms histone labeling. (**C**) SDS-PAGE analysis of Cy3-labeled FACT. (Left) Coomassie-stained gel showing the SPT16 and SSRP1 subunits and residual Sfp transferase used for labeling. (Right) Corresponding fluorescence scanning confirms SPT16 labeling.

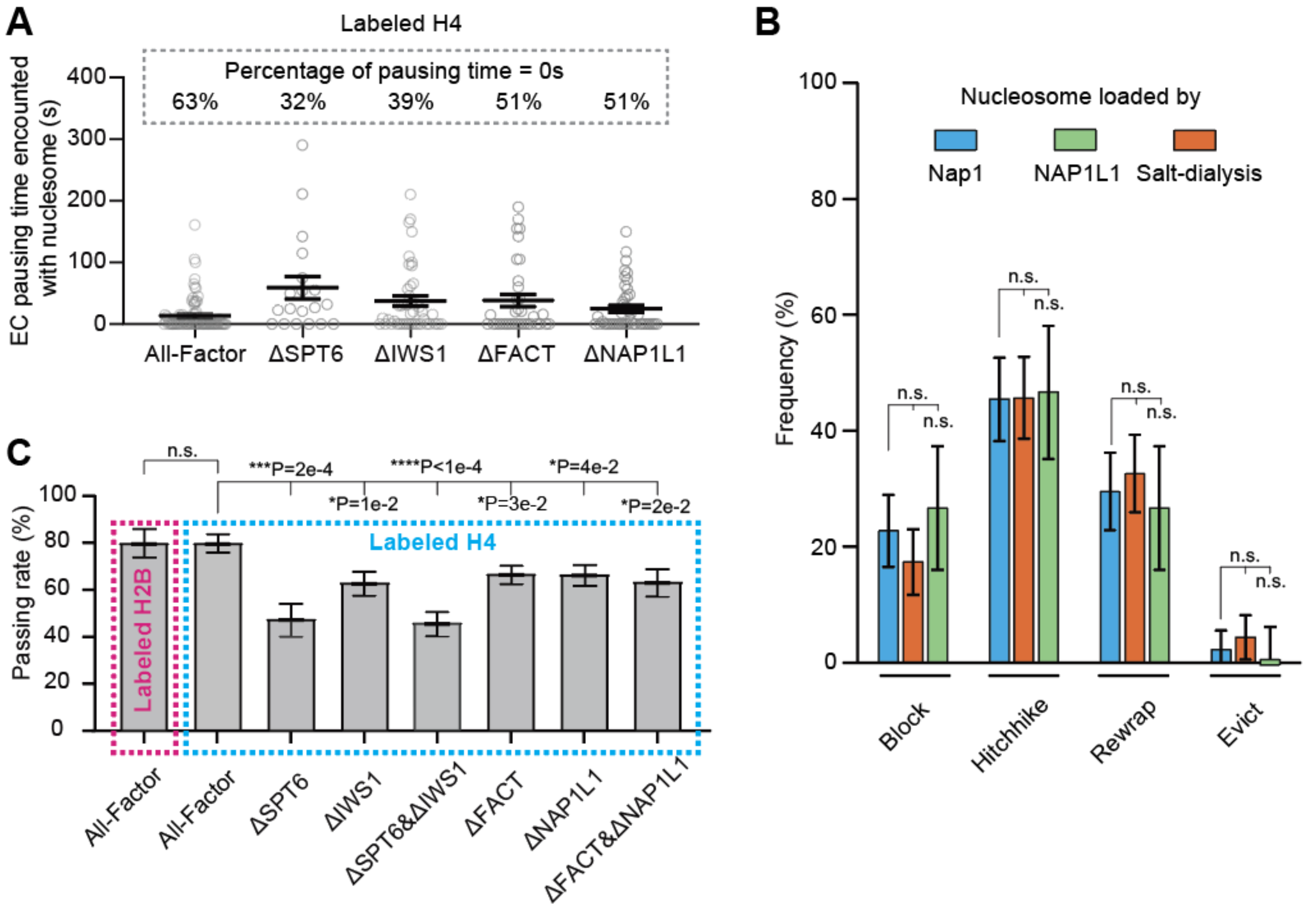


**Figure S2. Additional characterization of the outcome of EC-nucleosome encounter.**
(**A**) Distribution of the duration for which an EC paused before passing a nucleosome under the indicated factor conditions. Each point corresponds to an encounter event. Numbers above indicate the percentage of events without detectable pausing. (**B**) Frequency of different encounter outcomes for nucleosomes loaded by yeast Nap1, human NAP1L1, or salt dialysis. The outcomes are categorized based on the histone fate at the end of imaging. (**C**) Fraction of ECs that successfully passed a nucleosome using fluorescently labeled H2B or H4 and under the indicated factor conditions. Bars represent mean ± s.e.m. Statistical comparisons were performed using two-sided Fisher's exact tests.

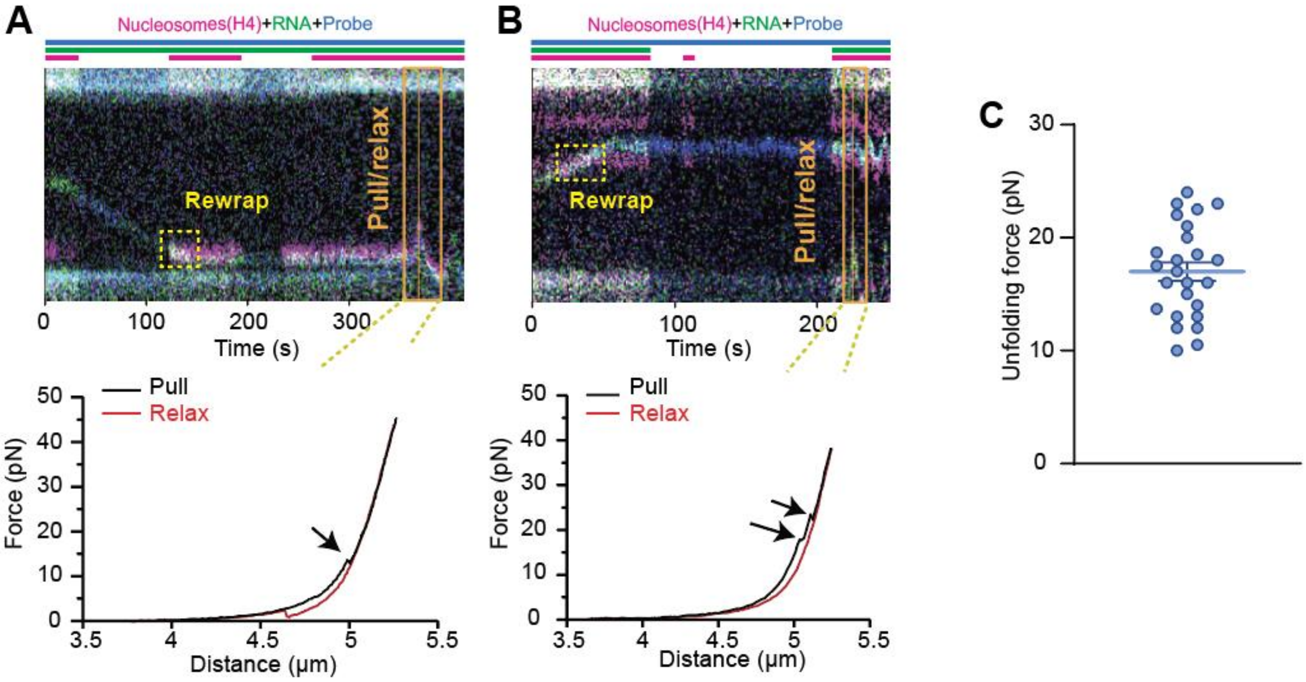


**Figure S3. Evaluation of nucleosome rewrapping after EC passage.**

(**A, B**) Representative data showing nucleosome rewrapping after EC passage. (Top) Kymograph showing the reappearance of nucleosome fluorescence signal (magenta; labeled at histone H4) after the traversal of EC (green and blue). Bars above indicate the time periods when specific lasers were switched on or off. (Bottom) Force-distance curve for the same tether undergoing a pull-relax cycle after imaging, which shows force-induced transitions characteristic of nucleosome inner-turn unfolding (arrows). (**C**) Distribution of the unfolding force from the force-distance curves. The values are consistent with previously reported measurements for nucleosome unwrapping. Bars represent mean ± s.e.m.

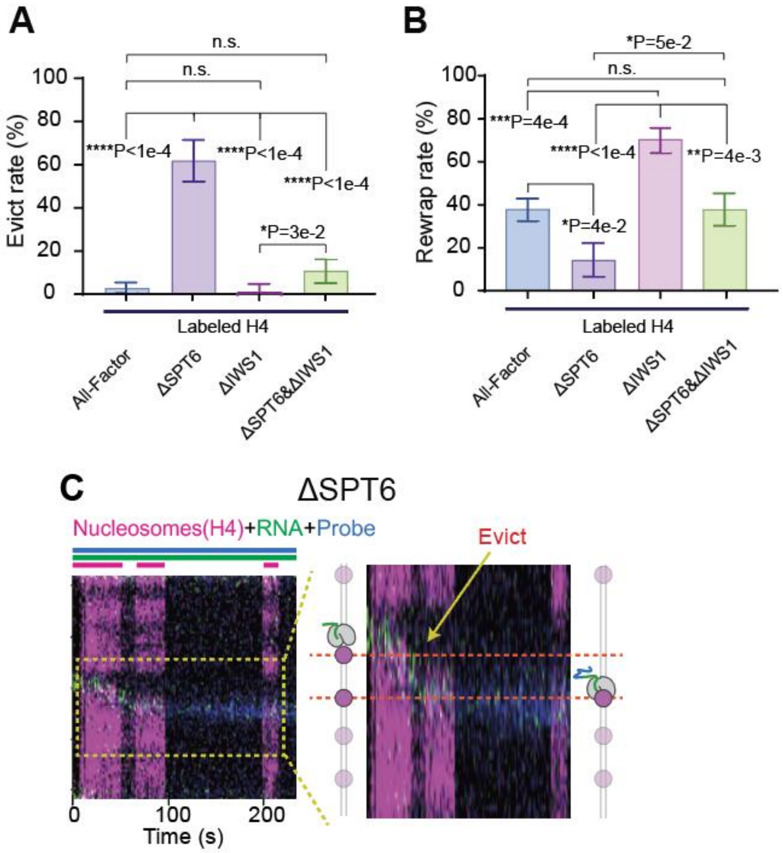


**Figure S4. Additional characterization of the effects of SPT6 and IWS1 omission on the histone fate after EC-nucleosome encounter.**
(**A**) Fraction of EC passage events that led to histone H4 eviction under the indicated factor conditions. (**B**) Fraction of EC passage events that led to H4 redeposition under the indicated factor conditions. The events are counted based on the histone fate at the end of imaging. Bars represent mean ± s.e.m. *P* values were calculated using two-sided Fisher's exact tests. (**C**) Representative kymograph for the ΔSPT6 condition showing eviction of H4 (loss of the magenta signal) after EC passage.

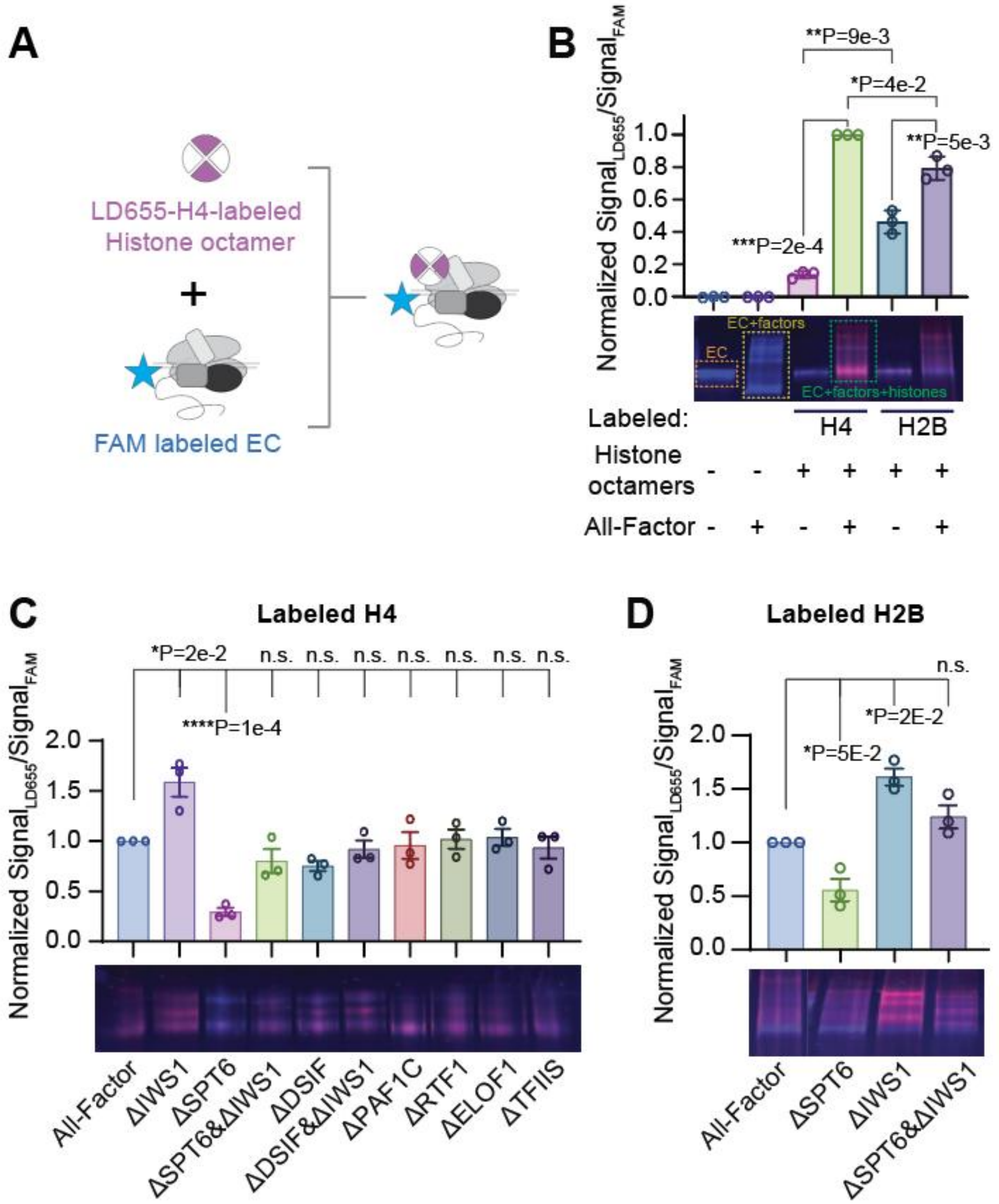


**Figure S5. A bulk assay for assessing EC-histone association.**
(**A**) Schematic of the experimental design. FAM-labeled EC without flanking DNA was incubated with LD655-labeled histone octamer and their association was assessed by EMSA on a native gel. (**B**) (Top) Quantification of the association (histone fluorescence signal normalized to the EC fluorescence signal). (Bottom) A representative gel showing the dependence of association on the presence of the octamer and elongation factors, which validates this assay. (**C, D**) Quantification of the EC-histone association when specific factors were omitted. The octamers were labeled at histone H4 in (C) or H2B in (D). The values are normalized to the All-Factor condition. Bars represent mean ± s.e.m. *P* values were calculated using unpaired two-tailed *t* tests with Welch's correction.

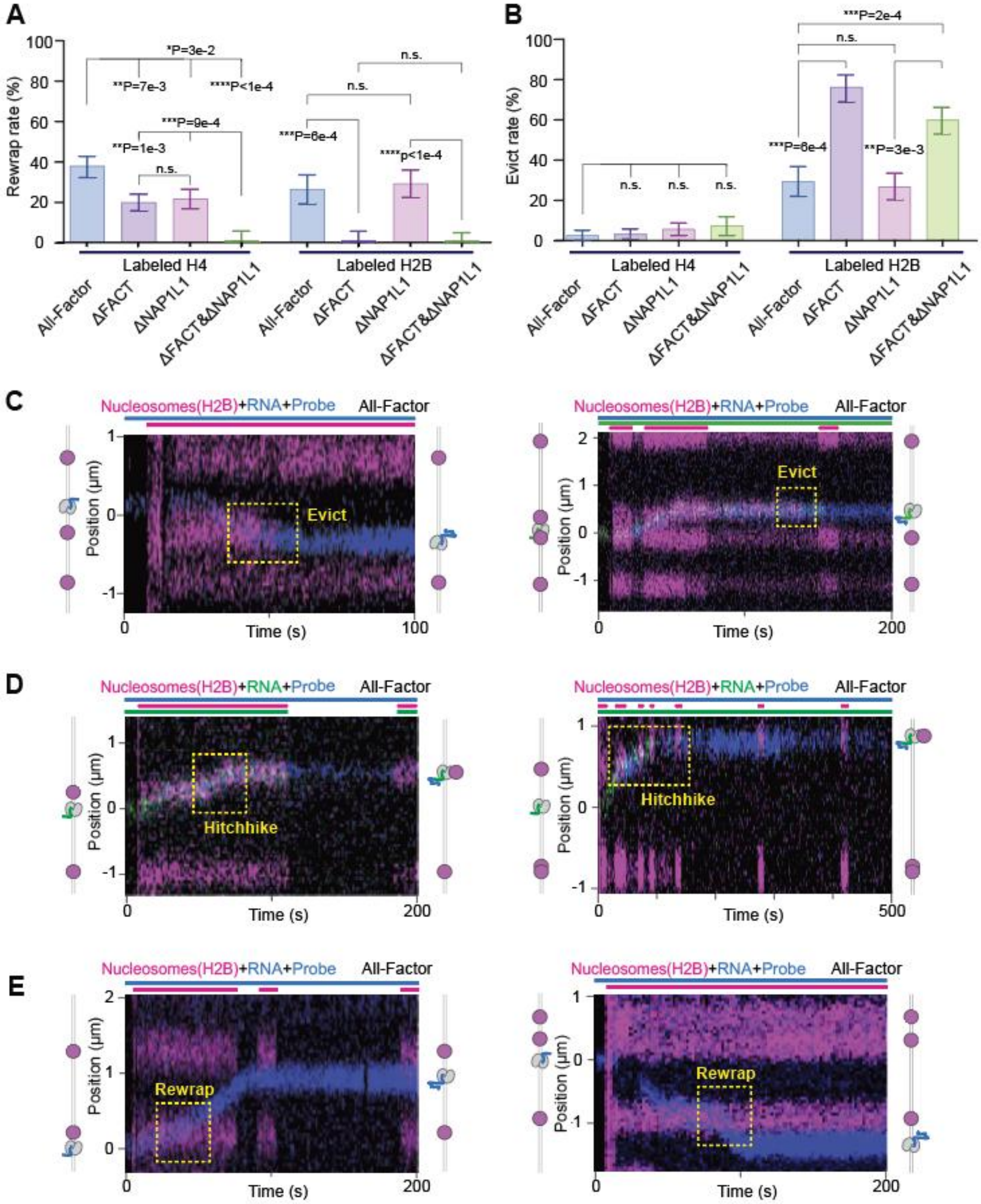


**Figure S6. Characterization of the histone H2B fate upon EC-nucleosome encounter.**
(**A, B**) Fraction of EC passage events that led to histone redeposition (A) or eviction (B) under the indicated chaperone omission conditions. The fates of histone H4 and H2B are both displayed. The events are counted based on the histone fate at the end of imaging. Bars represent mean ± s.e.m. *P* values were calculated using two-sided Fisher's exact tests. (**C–E**) Representative kymographs showing eviction (C), hitchhiking (D), and rewrapping (E) of H2B after an EC-nucleosome encounter under the All-Factor condition. Two examples are provided for each outcome. Bars above the kymographs indicate the time periods when specific lasers were switched on or off.

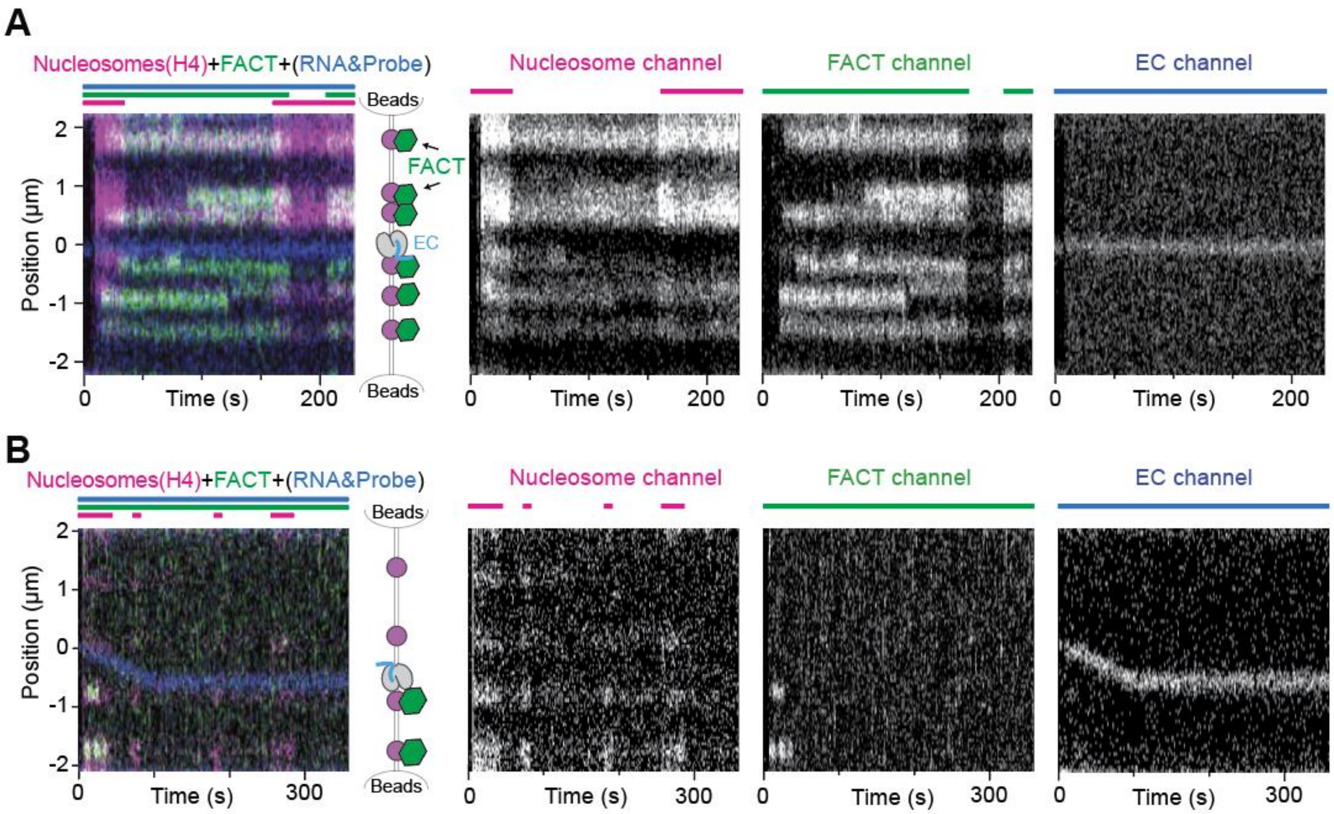


**Figure S7. Direct observation of FACT behavior during chromatin transcription.**
(**A**) Representative kymograph showing frequent FACT (green) colocalization with nucleosomes (magenta). (**B**) Representative kymograph showing no detectable FACT association with a translocating EC (blue) in the absence of histone hitchhiking. (Left) Overlaid fluorescence signals. (Right) Individual fluorescence channels. Bars above the kymographs indicate the time periods when specific lasers were switched on or off.

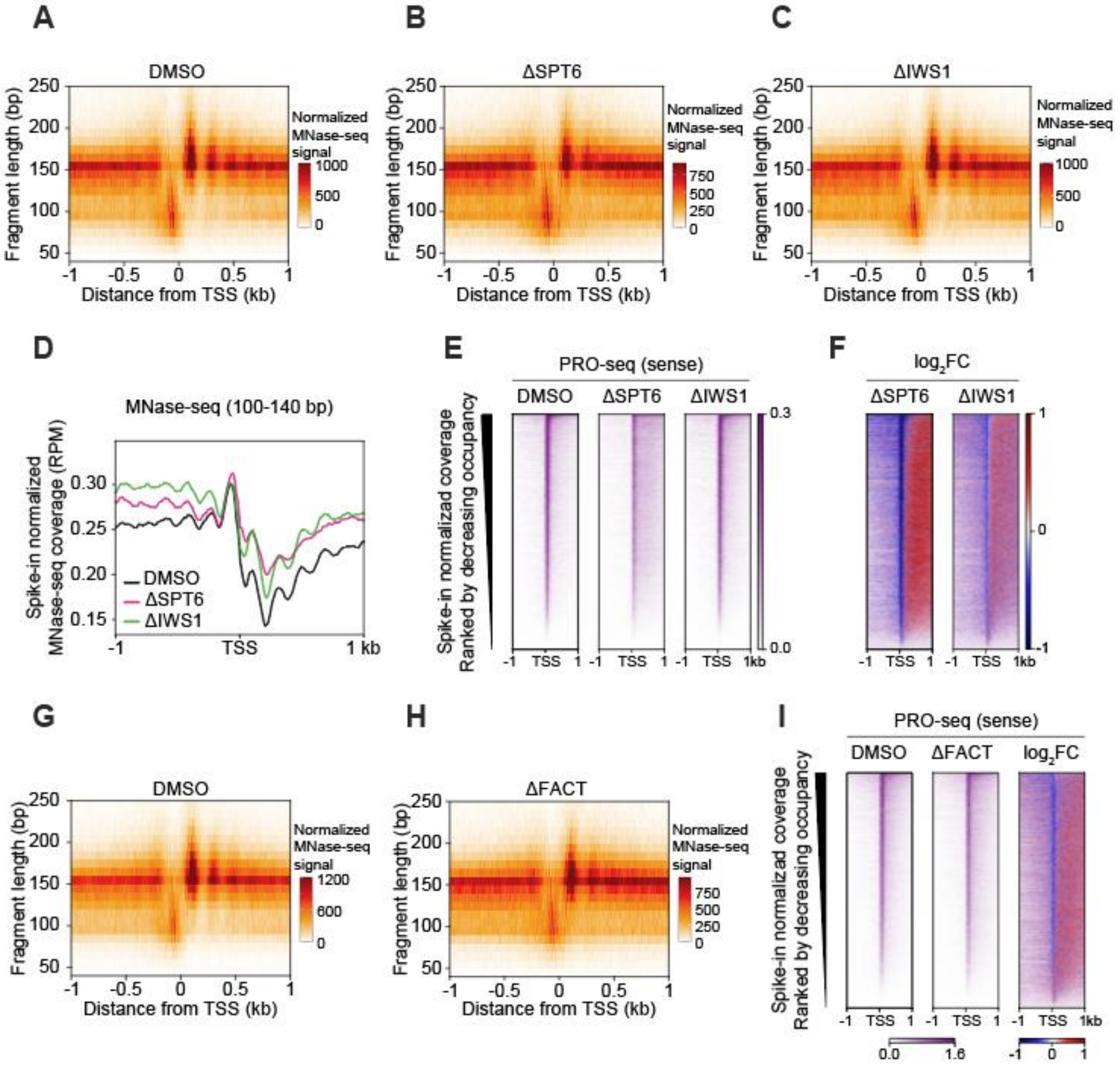


**Figure S8. Additional analysis of nucleosome organization and transcriptional activity in cells upon SPT6, IWS1, and FACT depletion.**

(**A–C**) V-plots showing library size–normalized MNase-seq signal as a function of fragment length and distance from TSSs in DMSO-treated control cells (A), SPT6-AID cells treated with IAA (500 µM, 3 h; ΔSPT6) (B), and IWS1-dTAG cells treated with dTAG-13 (200 nM, 3 h; ΔIWS1) (C). Fragment lengths from 50 to 250 bp and positions within ±1 kb of TSSs were binned at 10-bp resolution. (**D**) Metaplots showing the average spike-in–normalized MNase-seq coverage for 100–140-bp MNase-resistant fragments centered at TSSs under the conditions described in (A–C). (**E, F**) Heatmaps showing spike-in–normalized PRO-seq coverage for sense transcription centered at TSSs (E) and the corresponding $\log_2$ fold changes upon SPT6 or IWS1 depletion relative to DMSO-treated control cells (F). Genes are ranked by decreasing PRO-seq coverage in the DMSO-treated control, and the same gene order is used for all heatmaps. (**G, H**) V-plots showing library size–normalized MNase-seq signal as a function of fragment length and distance from TSSs in FACT-AID cells treated with DMSO (G) or IAA (1 mM, 6 h; ΔFACT)

(H). Fragment length and positional binning are the same as described in (A–C). (**I**) Heatmaps showing spike-in–normalized PRO-seq coverage for sense transcription centered at TSSs and the corresponding $log_2$ fold change upon FACT depletion relative to DMSO-treated control cells. Genes are ranked by decreasing PRO-seq coverage in the DMSO-treated control.

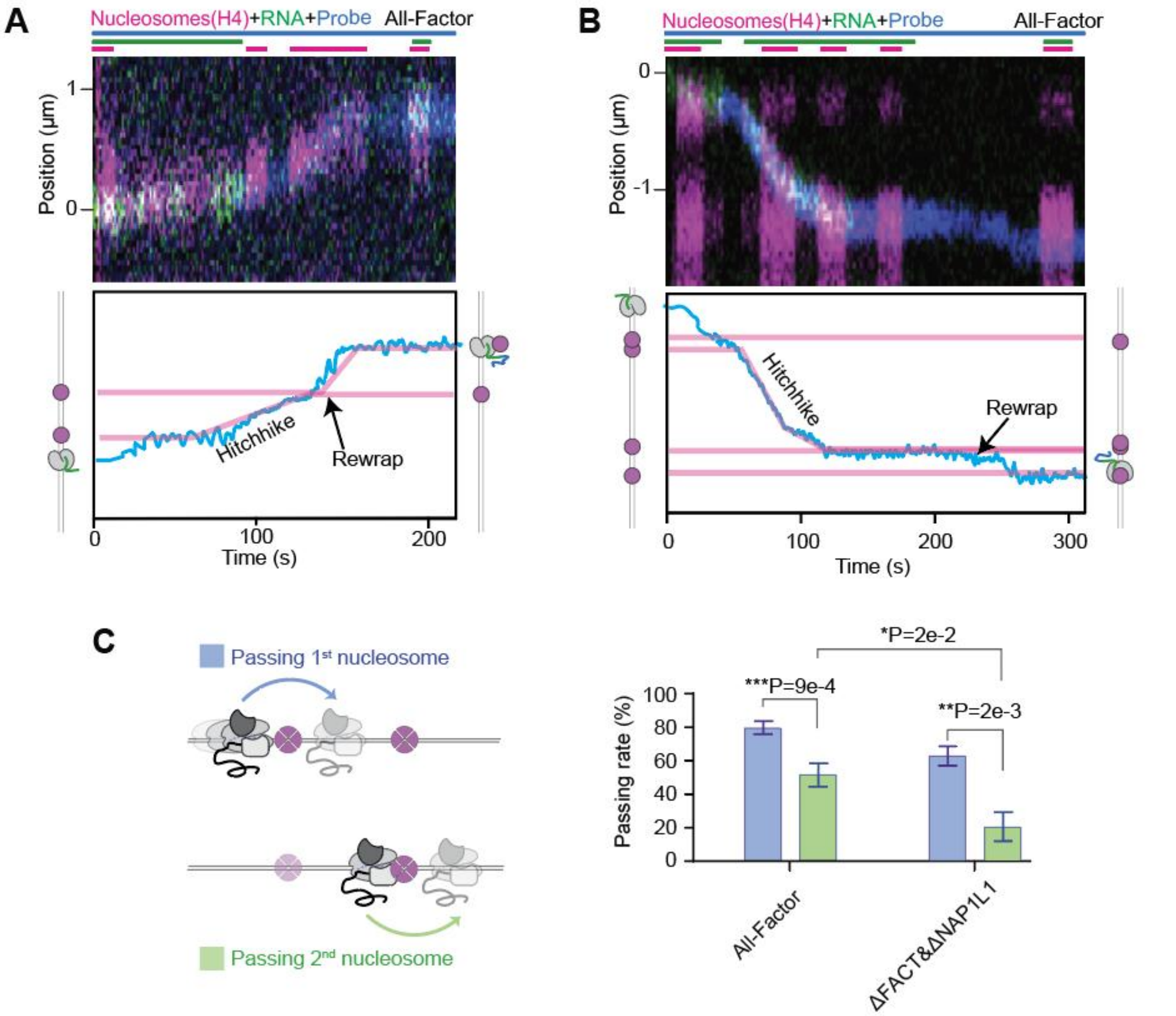


**Figure S9. Additional analysis of EC progression through successive nucleosomes.**
(**A, B**) Representative kymographs under the All-Factor condition showing a hitchhiking complex unloads its associated histones upon encountering a second nucleosome and then resumes translocation. Bars above the kymographs indicate the time periods when specific lasers were switched on or off. Plots below the kymographs show the extracted EC trajectory (cyan) and histone positions (magenta). (**C**) Fraction of ECs that successfully passed the first nucleosome (blue) and those that passed a second nucleosome (green) under All-Factor and ΔFACT&ΔNAP1L1 conditions. Bars represent mean ± s.e.m. *P* values were calculated using two-sided Fisher's exact tests.
